# Massive genome reduction occurred prior to the origin of coral algal symbionts

**DOI:** 10.1101/2023.03.24.534093

**Authors:** Sarah Shah, Katherine E. Dougan, Yibi Chen, Rosalyn Lo, Gemma Laird, Michael D. A. Fortuin, Subash K. Rai, Valentine Murigneux, Anthony J. Bellantuono, Mauricio Rodriguez-Lanetty, Debashish Bhattacharya, Cheong Xin Chan

## Abstract

Dinoflagellates in the Family Symbiodiniaceae (Order Suessiales) are diverse, predominantly symbiotic lineages that associate with taxa such as corals and jellyfish. Their ancestor is believed to have been free-living, and the establishment of symbiosis (i.e., symbiogenesis) is hypothesised to have occurred multiple times during Symbiodiniaceae evolution. Among Symbiodiniaceae taxa, the genus *Effrenium* is an early diverging, free-living lineage that is phylogenetically positioned between two robustly supported groups of genera within which symbiotic taxa have emerged. The lack of symbiogenesis in *Effrenium* suggests that the ancestral features of Symbiodiniaceae may have been retained in this lineage. Here we present *de novo* assembled genomes and associated transcriptome data from three isolates of *Effrenium voratum*. We compared the *Effrenium* genomes (1.2-1.9 Gbp in size) and gene features with those of 16 Symbiodiniaceae taxa and other outgroup dinoflagellates. Surprisingly, we find that genome reduction, which is often associated with a symbiotic lifestyle, predates the origin of Symbiodiniaceae. We postulate that adaptation to an extreme habitat (e.g., as in *Polarella glacialis*) or life in oligotrophic conditions resulted in the Suessiales ancestor having a haploid genome size ≤ 2Gbp, which was retained (or reduced) among all extant algae in this lineage. Nonetheless, our data reveal that the free-living lifestyle distinguishes *Effrenium* from symbiotic Symbiodiniaceae vis-à-vis their longer introns, more-extensive mRNA editing, fewer (∼30%) lineage-specific gene families, and lower (∼10%) level of pseudogenisation. These results demonstrate how genome reduction and the adaptation to symbiotic versus free-living lifestyles intersect, and have driven the diversification and genome evolution of Symbiodiniaceae.

## Introduction

Dinoflagellate algae in the Family Symbiodiniaceae comprise taxa that form symbioses with diverse marine organisms. Of particular importance to modern coral reefs, Symbiodiniaceae provide photosynthates *via* fixed carbon and essential nutrients to corals while resident in these cnidarians. The Symbiodiniaceae ancestor is believed to have been free-living (*1*) with members of this group forming symbiotic associations with corals as early as 230 million years ago (MYA) (*2*). Symbiogenesis, or the establishment of a symbiotic relationship between two or more taxa (*3*), can drastically influence lineage evolution, adaptation, and speciation as observed in obligate parasites and diverse symbiotic taxa (*4–8*). This phenomenon is termed the resident genome syndrome and was previously hypothesised to explain the observed patterns of Symbiodiniaceae genome evolution (*9*).

Based on current divergence time estimates for Symbiodiniaceae, the split between the basal genera (*Symbiodinium* and *Philozoon*) and the rest of the family occurred 166 MYA, whereas the more-recently branching symbiotic lineages diversified ∼109 MYA (*1*). If the emergence of symbiogenesis coincides with the earliest fossil evidence from 230 MYA (*2*), then different Symbiodiniaceae lineages would have arose and diversified during major global geological events. These events include the switch from aragonite to calcite seas (∼190 MYA; (*10*), the breakup of Pangea (150-230 MYA; (*11*), the diversification or extinction of potential hosts, e.g., the extinction of rudists 66 MYA (*12, 13*), and the overall change in coral morphology from the Triassic (201-252 MYA) to the Cretaceous (66-145 MYA) (*14, 15*). More-recent examples include the rapid radiation of the genus *Cladocopium* 4-6 MYA due to geographic isolation (*16*), and the co-diversification of *Symbiodinium fitti* and their coral hosts (*17*); see (*18*) for latest systematic revision of *Cladocopium* species.

As described above, symbiogenesis is expected to impact the genome evolution of symbionts to varying extents within a broad spectrum of “facultativeness” that reflects the nature of the host association (i.e., with obligate free-living and obligate symbiont at opposing extremes) and underpins evolutionary processes such as genome streamlining, genetic drift, expansion/contraction of mobile elements, pseudogenisation, gene loss, and varying mutation rates (*9*). Previous studies investigating the effects of symbiogenesis on Symbiodiniaceae genomes have focused almost entirely on symbiotic genera, with the polar-dwelling, highly specialised *Polarella glacialis* (*19*), a sister of Symbiodiniaceae and within the same Order Suessiales, providing the only free-living outgroup.

Haploid genome sizes of Symbiodiniaceae taxa and *P. glacialis* are estimated to be < 2 Gbp, based on available sequencing data (*19–22*), and < 5Gbp based on DNA staining and qPCR analysis of marker sequences (*23, 24*). Estimates based on DNA staining are generally larger than those based on sequencing data, likely due to the permanently condensed chromosomal structures of dinoflagellates that result in overestimation of DNA content (*25, 26*). The diverse dinoflagellate taxa external to the Symbiodiniaceae are predominantly free-living and, in comparison, have massive genome sizes, e.g., 4.8 Gbp estimated from sequencing data for the bloom-forming *Prorocentrum cordatum* (*27*), and 200 Gbp based on DNA staining for *Alexandrium tamarense* (*28*).

Among Symbiodiniaceae taxa, *Effrenium* is the early-diverging, exclusively free-living genus. The sole species, *E. voratum*, is globally distributed in temperate and subtropical waters (*1, 29*). Attempts to establish a symbiotic relationship between *E. voratum* and the anemone *Exaiptasia pallida* have been unsuccessful (*30, 31*). Current understanding of Symbiodiniaceae evolutionary history suggests that *E. voratum* diverged 147 MYA from the basal, largely symbiotic genera of *Symbiodinium* and *Philozoon*, and prior to the other later-diverging symbiotic genera (*1*). Whereas genomes of other free-living species such as *Symbiodinium natans* (*32*) have been generated, these taxa belong to genera that also include symbiotic species and thus might have experienced a symbiotic lifestyle at some point in their history. We expect the genus *Effrenium* to have remained unaffected by the influence of symbiogenesis, and thus retain the ancestral free-living lifestyle (and genome features) of Symbiodiniaceae.

In this study, we present *de novo* assembled genome and transcriptome data for three isolates of *E. voratum*. Incorporating publicly available genome-scale data from 16 Symbiodiniaceae taxa plus four free-living taxa external to the Symbiodiniaceae in a comparative genomic analysis, we examine genomic features in *E. voratum*. These include mobile elements, gene structures, gene-families, and pseudogenisation to gain insights into ancestral features of Symbiodiniaceae, and more broadly, Suessiales genome evolution.

## Results

### Genome-size reduction pre-dated divergence of Order Suessiales and Family Symbiodiniaceae

Using a mix of short- and long-read sequencing data (Table S1), we generated *de novo* genome assemblies for three isolates of *E. voratum* (assembly sizes 1.1-1.3 Gbp), with estimated haploid genome sizes of 1.2-1.9 Gbp (Table S2 and Fig. S1), completeness (BUSCO recovery 67.2-77.2%), and number of predicted genes (32,102–39,878) (Table 1, Tables S3 and S4) comparable to the other Symbiodiniaceae genomes (*19, 21, 22, 33-36*). We obtained all available genomic data from 23 dinoflagellate taxa: 19 from Symbiodiniaceae (Order Suessiales), two sister taxa of *P. glacialis* (Order Suessiales), and two distantly related free-living dinoflagellate taxa, *Prorocentrum cordatum* (Order Prorocentrales) and *Amphidinium gibbosum* (Order Amphidiniales). The 21 Suessiales taxa were grouped into: (a) the earlier-branching, largely symbiotic genus *Symbiodinium* (S1), (b) the three exclusively free-living *E. voratum* isolates (Ev), (c) the later-branching symbiotic Symbiodiniaceae lineages (S2), and (d) the free-living outgroup *P. glacialis* (Po) sister to Family Symbiodiniaceae (Table S3). The phylogenetic positions of these groups relative to other dinoflagellates are shown in Fig. 1A, along with light micrographs of representative species in S1, S2, and Ev (Fig. 1B). Cell size of *E. voratum* (12.2-13.3 µm (*29*)) is generally larger than S1 (e.g., *Symbiodinium microadriaticum* CassKB8; 8.0-11.0 µm (*23, 37*)) or S2 cells (e.g., *D. trenchii* CCMP2556; 7.5-10.0 µm (*23*)).

**Table 1.** Genome assemblies and gene predictions of *E. voratum* RCC1521, rt-383, and CCMP421.

| Isolate | RCC1521 | rt-383<br>(=CCMP3420) | CCMP421 |
| --- | --- | --- | --- |
| Location of isolation | Mediterranean<br>Sea off Blanes,<br>Spain | Eastern North<br>Pacific off<br>Santa Barbara,<br>USA | Cooks<br>Strait, New<br>Zealand |
| Reference | (29) | (133) | (134) |
| Genome sequencing technologies | Illumina,<br>PacBio,<br>Nanopore | Illumina,<br>PacBio,<br>Nanopore | 10X<br>Linked-reads |
| Genome assembly size (Gb) | 1.2 | 1.3 | 1.1 |
| Estimated genome size (Gb) | 1.4 | 1.2 | 1.9 |
| GC-content of genome assembly (%) | 50.8 | 50.6 | 50.9 |
| Total read coverage | 446× | 212× | 153× |
| Number of genome scaffolds | 3,881 | 11,607 | 38,022 |
| N50 of genome assembly (Kb) | 720 | 252 | 304 |
| Number of predicted genes | 32,108 | 39,878 | 32,615 |
| % BUSCO recovery (genome) | 34.0 | 35.1 | 20.4 |
| % BUSCO recovery (proteins) | 76.7 | 77.2 | 67.2 |

**Figure 1.**
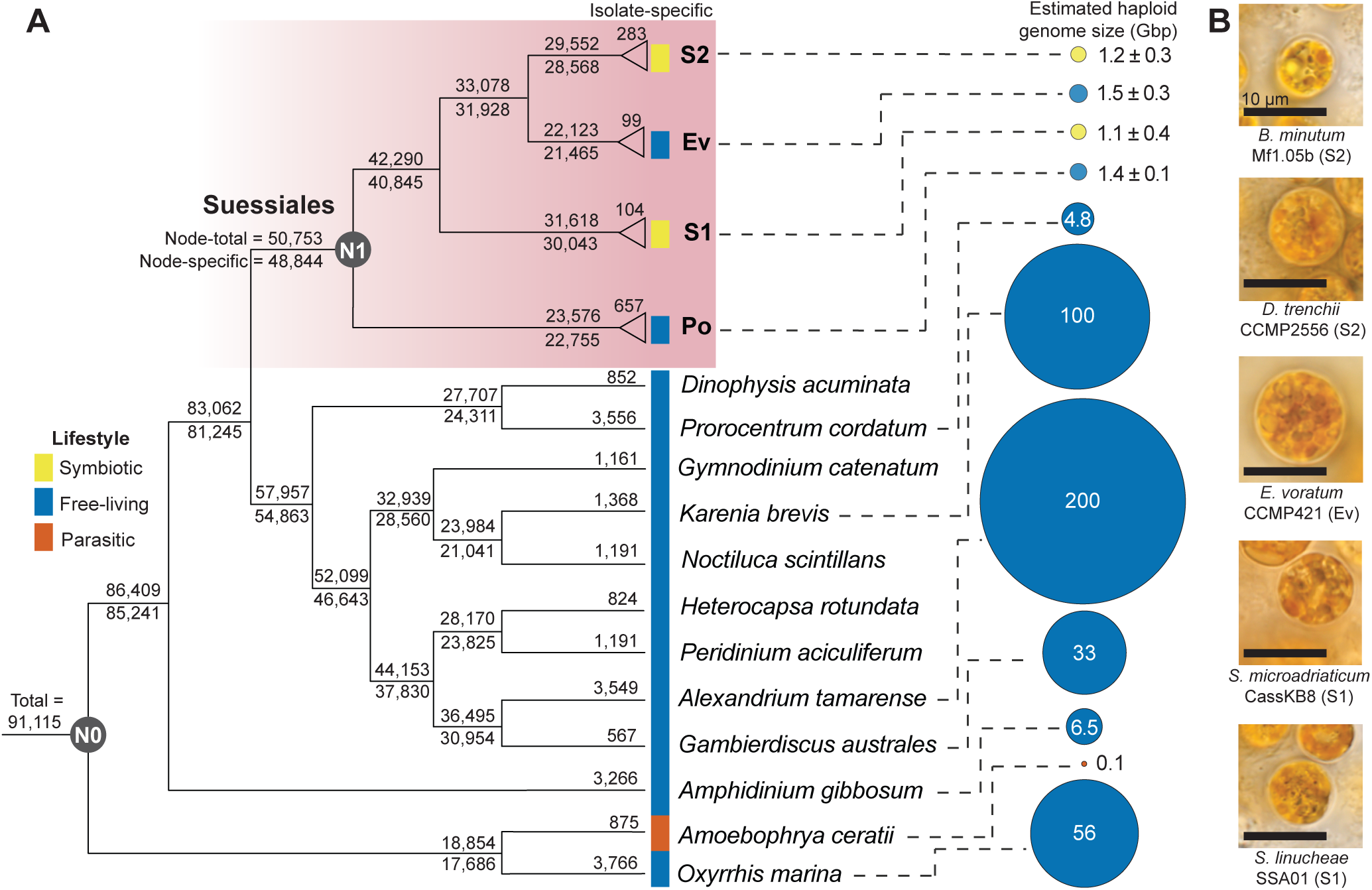
Species tree of dinoflagellates alongside estimated genome sizes. (**A**) The species tree was inferred using 91,115 orthologous protein sets derived from 1,420,328 protein sequences from 33 dinoflagellate taxa. Symbiotic lifestyles are shown in yellow, free-living in blue, and parasitic in orange. Node-total refers to the number of protein families which contain one or more taxa at the node. Node-specific are protein families that exclude taxa outside the node. Numbers at the tips of each branch represent isolate-specific protein families. For the S1, S2, Ev, and Po groups of Suessiales, the mean of isolate-specific protein families, and the mean ± standard deviation of estimated genome sizes are displayed. Node N0 contains homologous protein sets from all 33 taxa, whereas node N1 includes data from the 21 Suessiales. (**B**) Micrographs of representative taxa from the Symbiodiniaceae groups S1, S2, and Ev. Scale bar = 10 μm.

The most striking feature of genome evolution among Suessiales is the marked reduction in genome size that occurred in the common ancestor of this lineage. Whereas free-living dinoflagellates external to the Suessiales have genomes that range widely in size from ca. 5-200 Gbp, except the parasitic *Amoebophrya ceratii* that has a highly reduced genome of size 0.1 Gbp, all Suessiales genomes have a much narrower size range from 0.7-2.0 Gbp, estimated using sequencing data (Fig. 1A). This is accompanied by a ∼40% loss in gene families prior to the diversification of Suessiales, when compared to their closest dinoflagellate relatives (node N1 in Fig. 1A). This pattern is reminiscent of the red algae (Rhodophyta), whose common ancestor underwent massive genome reduction, precipitating the loss of canonical eukaryotic features such as flagella-based motility, phytochromes, and autophagy (*38*). These algae split into two monophyletic lineages, the extremophilic Cyanidiophytina that specialised to life in hot spring environments, and the species-rich mesophilic lineages (e.g., red seaweeds) that inhabit a variety of aquatic environments (*38*). Most red algae have therefore smaller genomes when compared to the green lineage and have adapted to diverse habitats through gene family evolution and horizontal gene transfer. In an analogous fashion in dinoflagellates, it appears that the Suessiales common ancestor underwent significant genome reduction, likely due to life in extreme habitats (e.g., the psychrophilic *Polarella glacialis*). This streamlining of the gene inventory may have played an important role in driving symbiotic associations with cnidarians that offered nutrient-rich and protected habitats within the animal tissues. The facultative lifestyle was likely retained in most Symbiodiniaceae because it offers the benefit of sexual reproduction during the free-living stage. The most substantial recovery from genome streamlining is offered by whole genome duplication, which has occurred in the *Durusdinium* lineage (*33*).

### Genome features of *E. voratum* versus early- and later-diverging symbiotic lineages

Genome sequences of the three *E. voratum* isolates share high similarity with 96% sequence identity over 93% of bases (Fig. 2A). Repetitive regions containing protein-coding genes were highly conserved relative those of other Suessiales (the Order containing Symbiodiniaceae and the earlier branching sister *P. glacialis*). We identified 98,344 core *k*-mers (*k* = 23; all possible 23-base sequences) that are common in genomes of all Suessiales taxa following Lo et al. (*39*), and recovered 95% of core 23-mers in repetitive regions of *E. voratum* (Fig. S2; see Materials and Methods). Among the three *E. voratum* genome datasets, we recovered 5-32 putative mitochondrial genome sequences in each dataset that encode the marker genes of *cob*, *cox1* and/or *cox3* (Table S5), and 6-32 putative plastid genome sequences in each dataset that encode the 16S/23S rRNA or one of the 11 plastid-encoded genes (*40*), including a putative empty minicircle sequence (Table S6).

**Figure 2.**
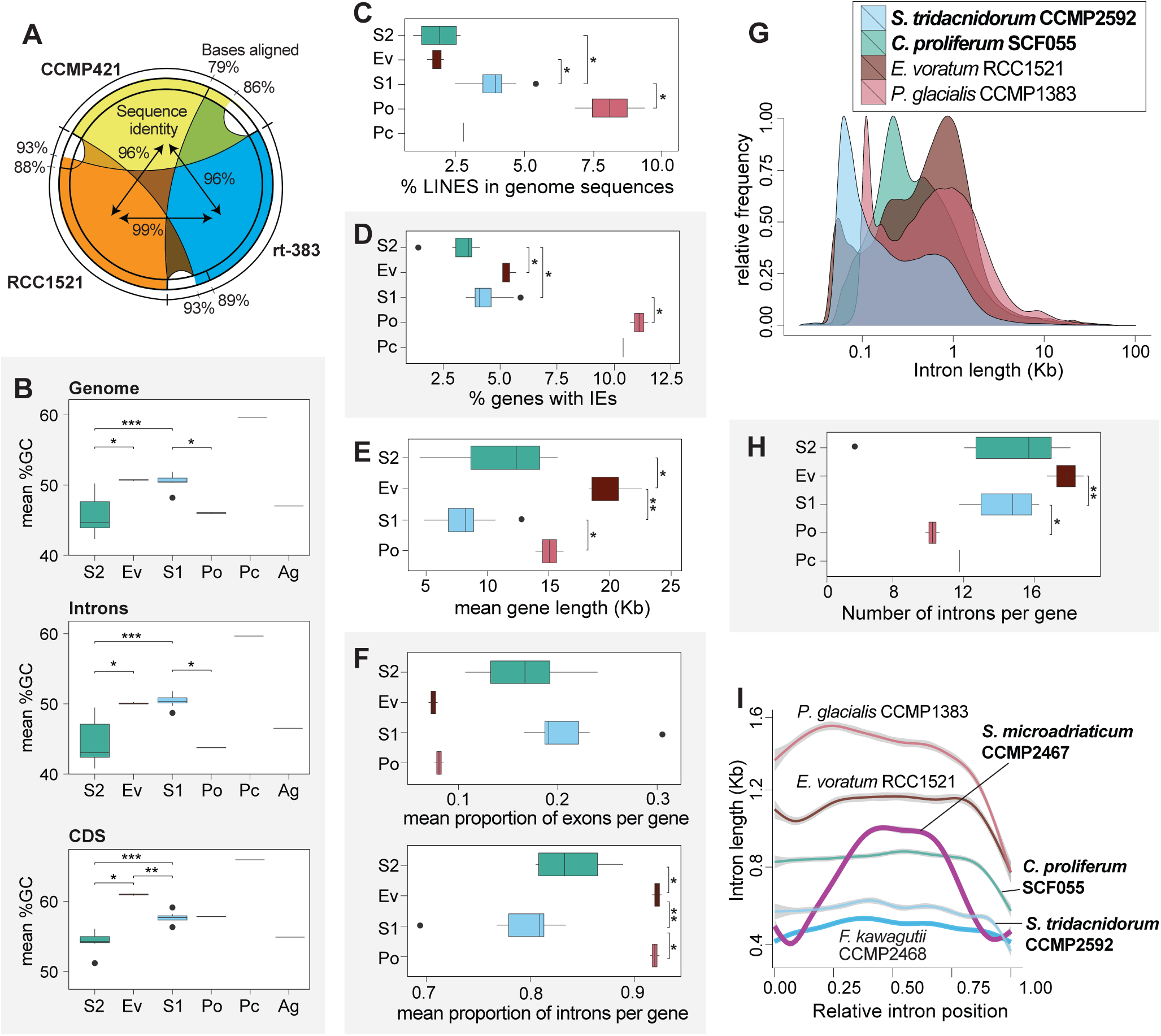
Genome features of *E. voratum* and other dinoflagellates. (**A**) Genome-sequence identity (italics within circle) and the percentage of aligned bases (outside circle) among the three *E. voratum* isolates. Features of representative genomes in the four Suessiales groups of S2, Ev, S1, and Po, plus *Prorocentrum cordatum* (Pc), in the order from the most-recent to most-ancient divergence, shown for (**B**) mean GC content in whole-genome, intronic, and CDS regions (*Amphidinium gibbosum* (Ag) was added for dinoflagellate-wide comparison), (**C**) percentage of mobile elements, and (**D**) percentage of genes containing introner elements (IEs). Gene features of Suessiales showing (**E**) mean gene length, (**F**) average proportions of exons and introns, (**G**) the relative frequency of introns by length (symbiotic lineages in boldface), (**H**) number of introns per gene, and (**I**) intron length versus relative intron position (symbiotic lineages in boldface). In all bar charts, *, **, *** represent *p* < 0.05, < 0.01, and < 0.001 respectively based on Wilcoxon rank sum test.

Given the history of genome reduction, we investigated the traits that may differentiate *Effrenium* from the symbiotic lineages of Symbiodiniaceae. We studied genome size, intron evolution, gene family evolution, pseudogenisation, RNA editing, and phylogenetic signals in Ev, S1 and S2, relative to the outgroup Po. The GC content of coding regions varied among the Symbiodiniaceae lineages (Fig. 2B) and were significantly lower in S2 (mean 54.2%; *p* < 0.05) relative to Ev (61.0%) and S1 (57.7%); in intronic regions, the mean GC is 44.6% (S2), 50.1% (Ev) and 50.4% (S1), whereas that of whole-genome sequences is 45.8% (S2), 50.7% (Ev), and 50.6% (S1). GC-rich genome sequences confer bendability to DNA helices (*41*) and may prevent cell freezing or desiccation (*42*). Variation of GC content in dinoflagellate genomes does not appear to correlate to lifestyle; among the free-living species external to Symbiodiniaceae, genomes of *P. glacialis* and *A. gibbosum* has a mean GC content of 46.4%, similar to S2, whereas the genome of *Pr. cordatum* has the highest GC content described thus far for any dinoflagellate, at 59.7% (*27*). Intracellular bacteria have a mutational bias towards low genomic GC content, e.g. ∼20% (*43*), but intracellular eukaryotes display both low and high extreme GC content patterns, ranging as low as 24% in the malaria parasite *Plasmodium falciparum* (*44*) to 67% in the green algal symbiont *Chlorella variabilis* in the ciliate *Paramecium* (*45*). The lower GC content in S2 genomes than the S1 counterparts underscores the dynamic nature of genomic GC content evolution in intracellular eukaryotes.

Mobile elements, particularly transposable elements (TEs), can influence genomic architecture and base composition, and have been used to reconstruct the evolutionary history of many species (*46–48*). Although facultative Symbiodiniaceae symbionts (i.e., S1 and S2) are expected to contain a larger proportion of mobile elements in their genomes when compared to free-living (Ev) lineages (*9*), no significant difference among the groups was observed in the overall abundance (Table S7 and Fig. S3A) or conservation of TEs (Fig. S3B). Although more contiguous genome assemblies generally contain more conserved repeats (Kimura substitution values centred around 3 for *D. trenchii* CCMP2556, *C. proliferum* SCF055 [formerly described as *C. goreaui* SCF055 (*18*)], the three *E. voratum* isolates*, S. natans* CCMP2548, and *S. tridacnidorum* CCMP2592) compared to the others (i.e., values centred around 20), we note diverged repeats (Kimura values centred around 25) in the chromosome-scale assembly of *S. microadriaticum* CCMP2467 (Fig. S3B). This result suggests a potential technical bias in the recovery of mobile elements. Despite this issue, based on the proportions for distinct types of mobile elements in each group, we found significantly (*p* < 0.05) more long interspersed nuclear elements (LINEs) in S1 (3.9%) compared to Ev (1.8%) and to S2 (1.9%), with the proportion of the outgroup Po at 8% (Fig. 2C). This result suggests that S1 likely retains ancestral LINEs, lending support to the notion of loss of LINEs after the diversification of basal Symbiodiniaceae genera (*21*).

Introner elements (IEs) are a type of mobile element consisting of inverted and direct repeat motifs found at 5′- and 3′-end of introns in diverse eukaryotes (*49–52*). Recent research on dinoflagellates revealed that IEs are more abundant in free-living (within 10-12% of genes) than in symbiotic/parasitic species (0.8-6.0% of genes) (*27, 53*). Although not as high as in other free-living dinoflagellates, the Ev genomes exhibit more IE-containing genes (5%) than do the S1 (4%) and S2 genomes (3%) (Fig. 2D). IEs have been postulated to be non-autonomous and their mobility is dependent on transposases encoded in dinoflagellate genomes (*53*). We recovered transposase protein sequences from most of the Symbiodiniaceae genomes in this study (Table S8), suggesting a capacity for IEs to be mobile.

Significantly (*p* < 0.05) longer genes were observed in Ev (mean 20 Kb) than S1 (8 Kb) and S2 (11 Kb) (Fig. 2E), primarily driven by longer intron sizes (introns make up, on average, 92% of a gene [Fig. 2F]; sizes peak at 1 Kb [Fig. 2G]) and higher intron density per gene (mean of 18 for Ev, 14 for S1, 14 for S2) (Fig. 2H). The driving mechanism for this trend may reflect one of two evolutionary scenarios: (a) intron expansion in Ev, or (b) intron contraction in S1/S2. In examples of the first scenario, TE-mediated insertions drive intron expansion and are biased toward the 5′ end of genes to prevent disruption of functional elements (*54*), yielding larger intron sizes at 5′ ends. We did not observe this trend in Ev (Fig. 2I). In the second scenario, which has been observed in endosymbiotic/parasitic organisms, the reduction of intron size and density occurs as a result of genome reduction and/or streamlining induced by spatial confinement in the host organism or cell (*46, 55*).

Considering the evolutionary history of Symbiodiniaceae, intron contraction in S1/S2 taxa due to their symbiotic lifestyle is more plausible than intron expansion in Ev. Large introns observed in Po (mean 1.4 Kb) (Fig. 2G), albeit at a lower intron density per gene (Fig. 2H), lend further support to this notion.

### Symbiogenesis shaped evolution of gene families and post-transcriptional processing in Symbiodiniaceae

To examine the effect of symbiotic lifestyle on protein family evolution in Symbiodiniaceae, we first inferred 53,173 homologous protein families from all 811,611 protein sequences predicted from the 21 Suessiales genomes (see Materials and Methods). Most protein families (47,353 of 53,173 [89%]) were shared among the four groups (S1+S2+Ev+Po). With respect to functions annotated in all families, these families were enriched in functions such as cellular motility, biosynthetic processes for rRNA, antibiotics, and glycosides (Table S9).

There were more lineage-specific protein families in S1 (6,389) and S2 (4,056) than in Ev (1,734), and the two symbiotic groups (S1+S2) shared 3,357 protein families not found in the other groups (Fig. 3). The high number of protein families present only in symbiotic lineages that split from each other over 40 million years of evolution suggests convergent evolution due to the symbiotic lifestyle. These protein families were enriched for diverse functions including signalling, apoptosis, protein splicing, photosynthesis, cell adhesion, and various transferase activities (Fig. 3). Incidentally, Po shared more protein families with symbiotic lineages (1,290; S1+S2+Po) than with Ev (221; Ev+Po); these families were enriched in functions such as autophagy and microtubule organisation.

**Figure 3.**
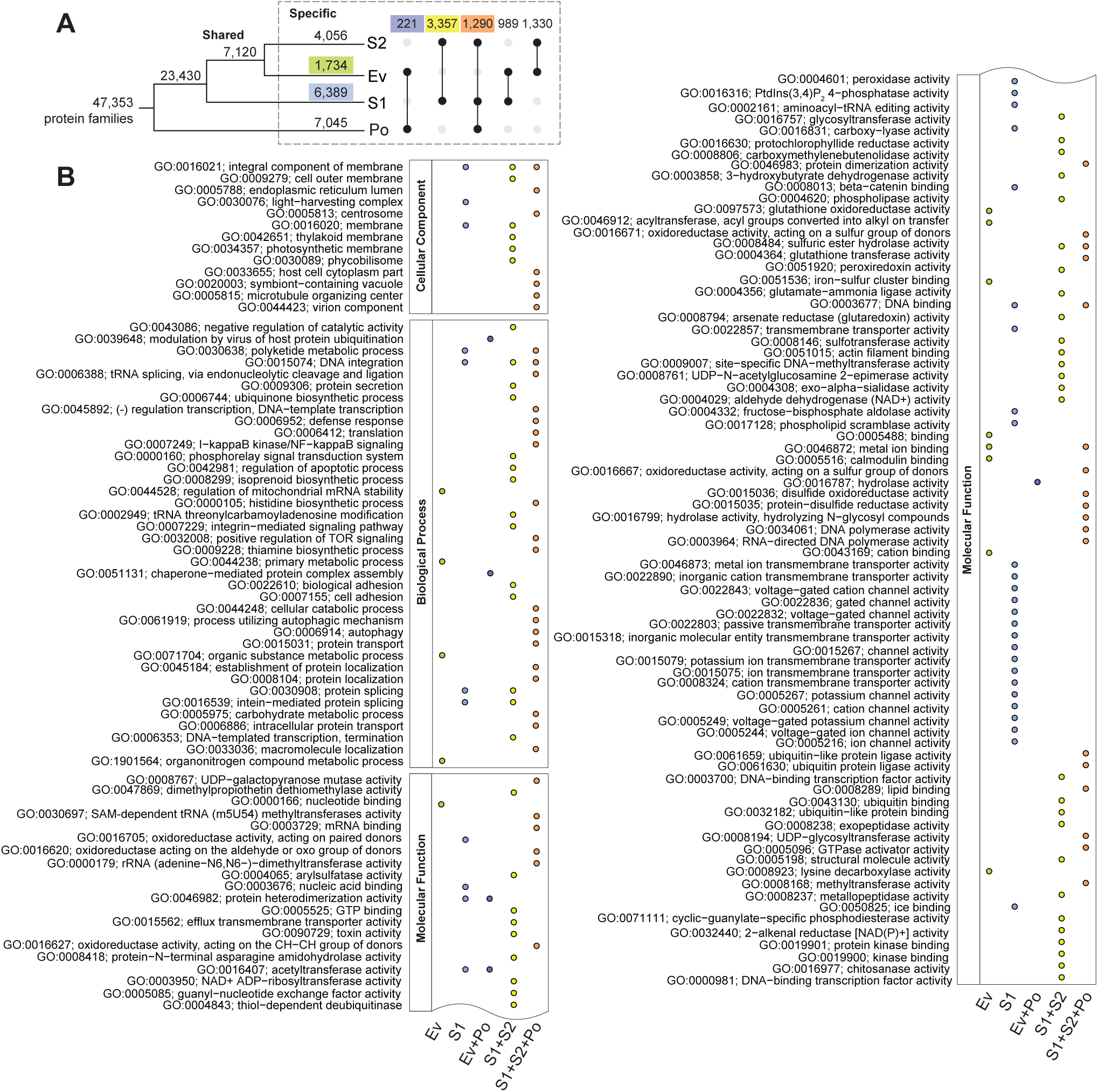
Gene family evolution in Suessiales. (**A**) Number of protein families is shown at each node and branch represents those that are shared among or specific to S1, Ev, S2, and/or Po. Number of families that are exclusive to Ev (green), to S1 (light blue), to Ev+Po (dark blue), to S2+S1 (yellow), and to S2+S1+Po (orange) were highlighted. (**B**) Enriched Gene Ontology (GO) terms for genes in the five distinct groups relative to all GO terms in the corresponding taxa, arranged in decreasing order of significance from top to bottom within the categories: Cellular Component, Biological Process, and Molecular Function.

Facultative symbionts in Symbiodiniaceae are expected to display higher levels of pseudogenisation, a major feature of the resident genome syndrome (*9*). To investigate this issue, we identified putative pseudogenes in the Symbiodiniaceae genomes (Table S10) following González-Pech et al. (*21*). We defined the level of pseudogenisation, ψ, as a ratio of the number of putative pseudogenes to the number of putative functional genes per homologous family; see Materials and Methods. We compared ψ independently for Ev (ψ_Ev_) against that for S1 (ψ_S1_), S2 (ψ_S2_), and the combined S1 and S2 (ψ_S1+S2_), then identified protein families that exhibited significant difference (*p* < 0.05) of this value. More protein families display ψ_S1_ > ψ_Ev_ (336) and ψ_S2_ > ψ_Ev_ (273; Fig. 4A), compared to ψ_S1_ < ψ_Ev_ (300) and ψ_S2_ < ψ_Ev_ (126; Fig. 4B). There was nine-fold more protein families exhibiting ψ_S1+S2_ > ψ_Ev_ (229; Fig. 4A) than vice versa (25; Fig. 4B). These pseudogenes are associated with a wide range of functions, including cell cycle processes and stimuli response (Figures 4C and 4D). The protein families that display significantly higher ψ in the symbiotic lineages are mostly mutually exclusive from the 3,357 families that putatively experienced convergent evolution (only 16–23 families are represented in ψ_S1_, ψ_S2_, ψ_S1+S2_). We found negligible technical biases in the clustering of homologous sequences that may affect our inference of pseudogenes, i.e., protein families displaying ψ remained stable at different clustering parameters (Fig. S4; see Materials and Methods). These results suggest that in addition to convergent evolution in the symbiotic lineages, these lineages of Symbiodiniaceae have experienced a greater extent of pseudogenisation than has the free-living Ev.

**Figure 4.**
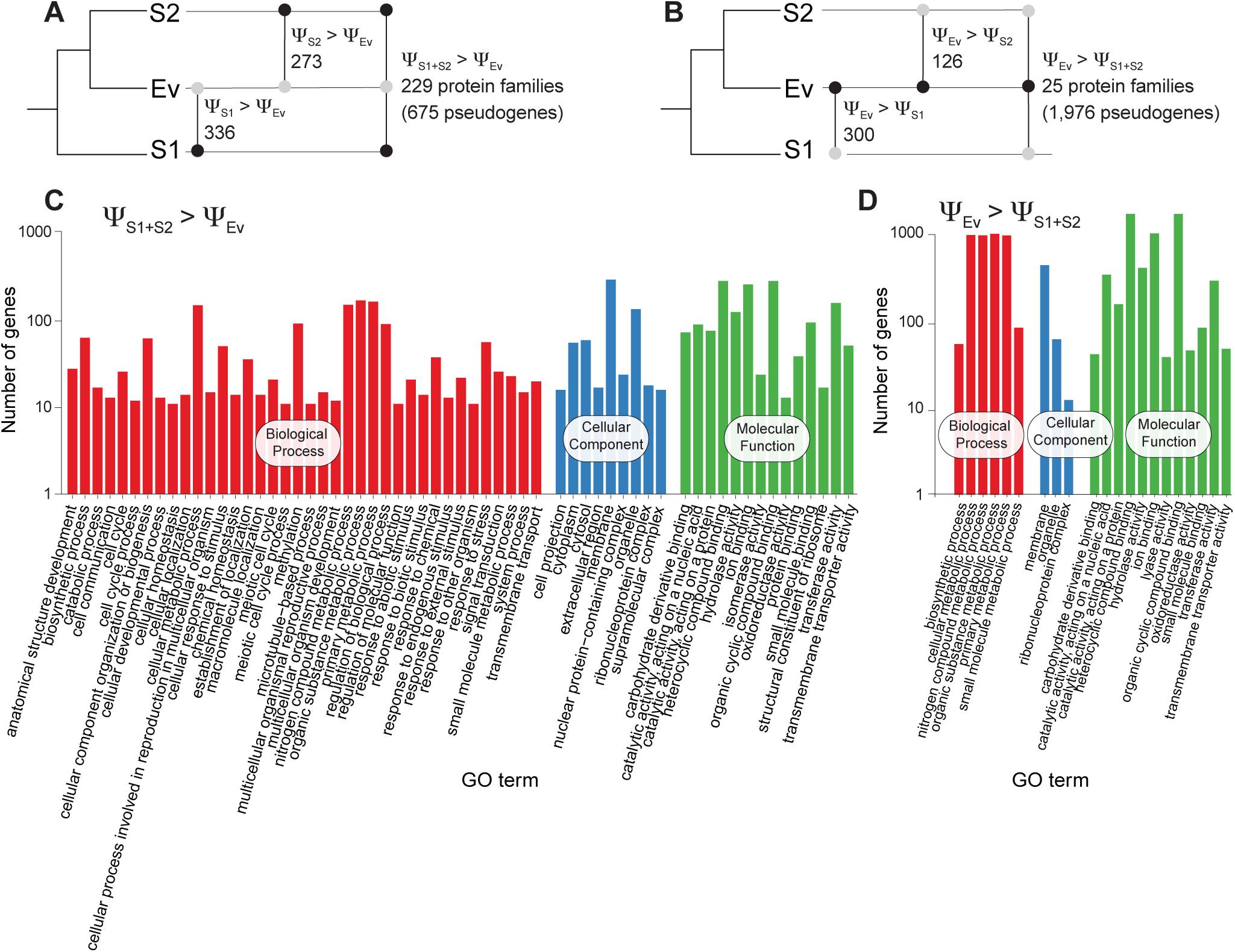
Pseudogenisation in Symbiodiniaceae. Number of protein families with significantly different level of pseudogenisation, ψ, shown for (**A**) those with greater extent in symbiotic lineages (i.e., higher ψ in S1, S2, or S1+S2, relative to Ev), and (**B**) those with greater extent in the free-living Ev (i.e., higher ψ in Ev relative to S1, S2, or S1+S2). The black circles on the upset plots indicate taxa groups with higher ψ than those with grey circles. The associated GO terms are shown for (**C**) those where ψS1+S2 > ψEv, and (**D**) those where ψEv > ψS1+S2.

Editing of mRNAs allows Symbiodiniaceae to increase the variability in protein isoforms (*56*). To assess mRNA editing in *E. voratum*, we focused on RCC1521, which has the most contiguous genome assembly, and compared the rates and types of mRNA editing against a representative from S1 (*S. microadriaticum* CCMP2467 (*56*)) and S2 (*D. trenchii* CCMP2556 (*33*)). We identified 45,009 unique mRNA edited sites in *E. voratum*, about 13-fold and 4-fold greater than in *S. microadriaticum* and *D. trenchii*, respectively (Table S11). A larger proportion of genes in *E. voratum* (9,158, 28.5%) contain mRNA edits, compared to *S. microadriaticum* (774, 1.6%) and *D. trenchii* (4,227, 7.6%), although the distribution of substitution types is similar among the three species (Fig. S5A-D). Most edits in *D. trenchii* and *E. voratum* (> 60%) are located in exons (Fig. S5E), whereas in *S. microadriaticum* edits are evenly split between exons and introns, although this may be due to the more-fragmented genome assembly of Liew et al. (*56*). As also observed in *S. microadriaticum* (*56*), there is a slight bias for edits to occur near 5′ ends of genes (within the first ∼10% of gene length, *p* < 0.05; Fig. S5F) and the edits tend to be located within 1 Kb of each other (versus the distribution expected at random, *p* < 0.05; Fig. S5G). The high level of mRNA editing in Ev is consistent with data from another free-living dinoflagellate, *Pr. cordatum* (42,969 edited sites; 32,067 within 10,169 [12%] genes) (*27*). Together with our observation of fewer Ev-specific protein families than those specific to S1 or S2 (Fig. 2), these results suggest a more pronounced role of mRNA editing in generating functional diversity in free-living versus symbiotic dinoflagellate taxa. The relationship and impact of this RNA editing on the encoded proteins remain to be investigated using proteomics.

### Impact of symbiogenesis on phylogenetic signal of non-coding regions

Our alignment-based phylogenetic trees inferred using multiple protein families (Fig. 1A), the standard molecular marker of 18S rDNA (Fig. 5A), and the ITS2 (Fig. S6A) were consistent with previously established phylogenies (*1, 57, 58*). We then analysed the phylogenetic signal of whole-genome sequence data using a *k-*mer-based alignment-free (AF) approach (see Materials and Methods), focusing on distinct genome-sequence regions following Lo et al. (*39*). Interestingly, the inferred AF phylogenies of introns (Fig. 5B), repetitive regions (Fig. S6B), repeat-masked whole-genome sequences (Fig. 5C), and entire whole-genome sequences (Fig. S6C) placed Ev as the basal group, branching earlier than S1/S2. In comparison, the AF phylogenies for coding regions (i.e., coding sequences [CDS; Fig. 5D] and protein sequences [Fig. S6D]) were largely congruent with the 18S rDNA phylogeny (Fig. 5A), placing S1 as earlier branching than Ev. The branching orders of Ev vs S1/S2 in AF trees were supported by a robust jackknife support of ≥ 96% based on 100 subsampled replicates (Fig. 5B-D; see Materials and Methods). This trend is consistent with visualisation of the AF distances as a phylogenomic network (Fig. S6E), in which *E. voratum* is more closely related to *P. glacialis* based on intron sequences, and to *Symbiodinium* spp. based on CDS.

**Figure 5.**
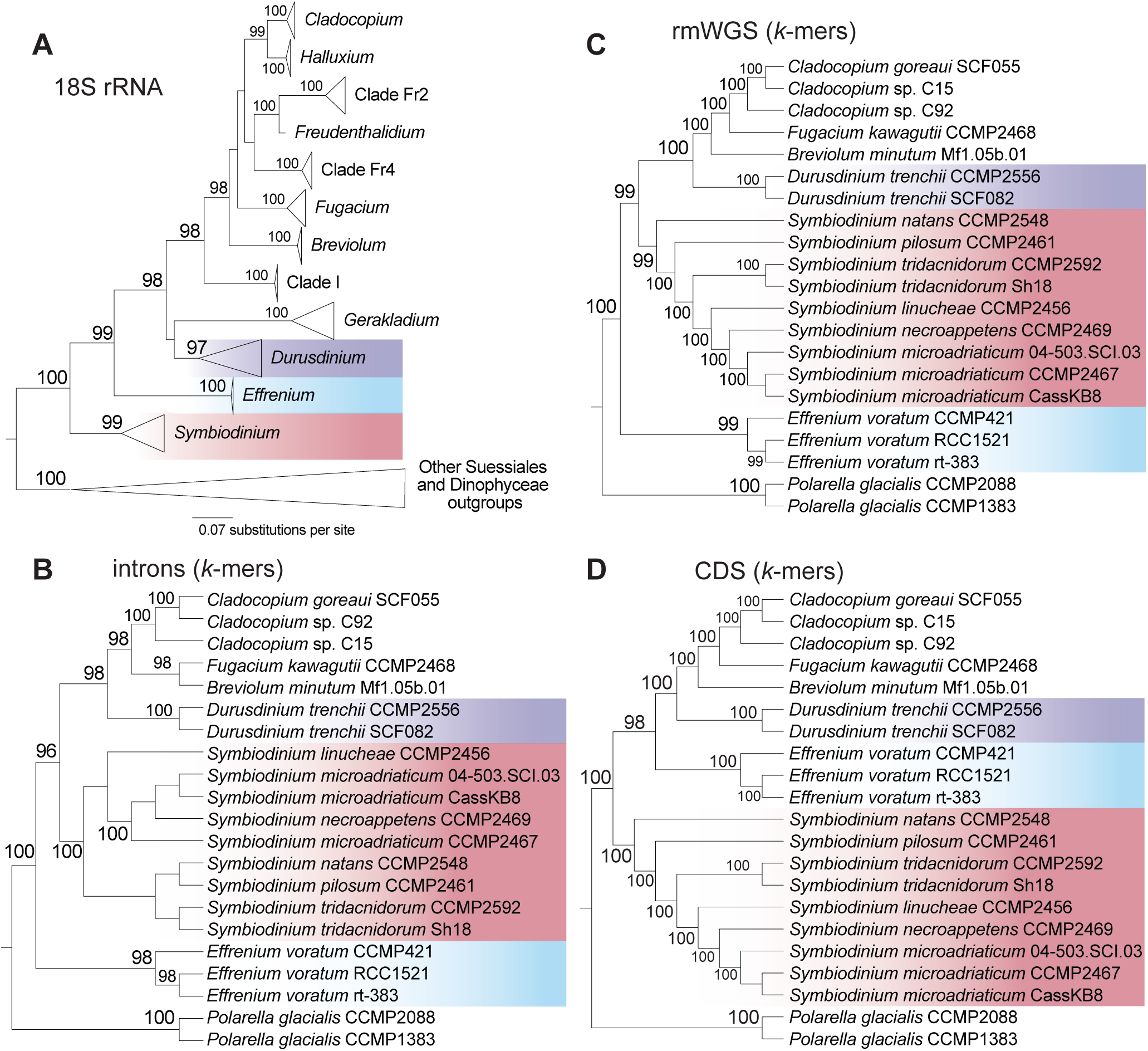
Phylogenetic relationship of *E. voratum* within Symbiodiniaceae and other taxa. (**A**) Maximum likelihood species tree of dinoflagellates inferred based on alignment of 18S rDNA sequences from 13 representative taxa, showing node support derived from ultrafast bootstrap of 2,000 samples; unit of branch length is number of substitutions per site. Tree topologies inferred from *k*-mers in 21 Suessiales genomes using our alignment-free (AF) approach are shown for (**B**) intronic regions (*k* = 21), (**C**) repeat-masked whole-genome sequences (k = 23), and (**D**) coding sequence regions (*k* = 19). Jackknife support of _ 96% based on 100 pseudo-replicates is shown on each node of the AF trees.

The incongruence between phylogenies of coding and non-coding regions with robust support of the distinct clades clearly indicate differential selective pressure acting on these two regions in Symbiodiniaceae genomes, as demonstrated in an earlier study (*39*). This is likely explained by incomplete lineage sorting, horizontal gene transfer, hybridisation, or convergent GC-biased gene conversion (*59–61*). This result may also reflect the retention of ancestral non-coding regions in the *E. voratum* genomes, and/or the loss of some non-coding regions in symbiotic lineages due to genome streamlining.

### Symbiogenesis differentially affected early-versus late-branching symbiotic lineages of Symbiodiniaceae

Following a possible genome reduction in the free-living ancestor of the Suessiales, some lineages may have developed symbiogenesis with a range of hosts, giving rise to the Symbiodiniaceae Family. Common and distinct genomic features we observed between early and late-branching symbiotic lineages of Symbiodiniaceae suggest an interplay between the geological eras during which they arose, and the corresponding coral morphology and ocean chemistry (Fig. 6). Ancestral Symbiodiniaceae inhabited stony corals presumably as early as 230 MYA in the late Triassic (*2*) and may have driven the Norian-Rhaetian reef bloom (*62*). These early Scleractinian corals (e.g., *Retiophyllia*) tended to be uniserial, i.e., possessing one corallite per branch, and phaceloid with thick walls (*63, 64*), and thus were less efficient at harvesting light (*65*). The ability of the early-diverging extant *Symbiodinium* to thrive under high or variable light (*1*) may be a trait inherited from their ancestor living in these ancient corals. Because these early Symbiodiniaceae adapted to different hosts, they likely underwent genome streamlining (*9*), experienced high pseudogenisation and a reduction in mRNA editing and intron sizes (Fig. 6), as observed here and in other studies (*21, 56*). Although these trends were also observed in the later-branching symbionts, *Symbiodinium* uniquely retained ancestral LINE repeats (Fig. 2C) which were lost in later-branching Symbiodiniaceae including *E. voratum*.

**Figure 6.**
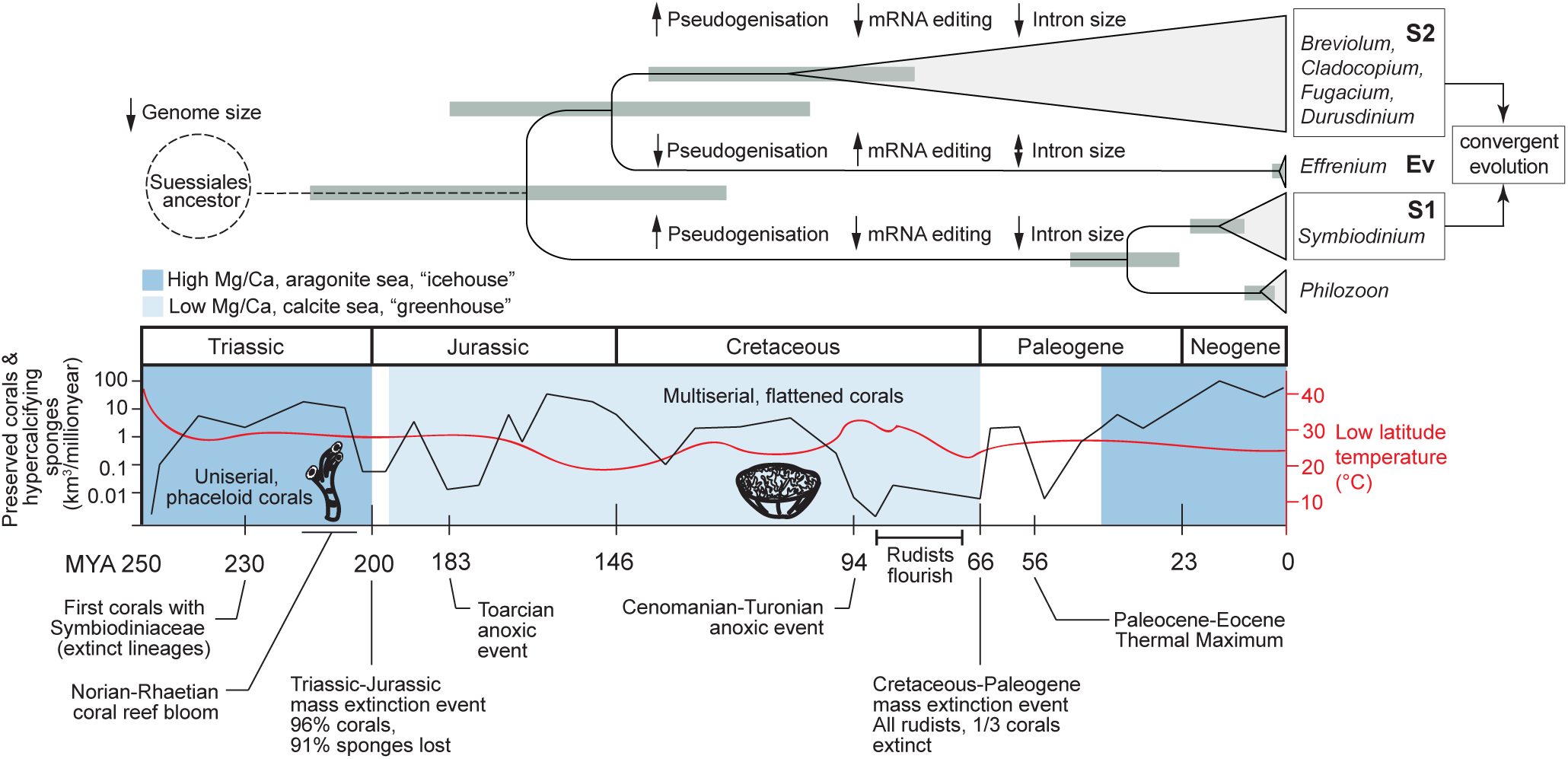
Timeline of Symbiodiniaceae genome evolution and coral evolution. The estimated divergence timeline of the Family Symbiodiniaceae (*1*) is shown at the top, indicating representative taxa of S1, Ev, and S2. Grey bars represent 95% confidence intervals of divergence times. Key genome signatures for each group related to pseudogenisation, mRNA editing, and intron size are shown along the branch. The dotted line represents the yet-unknown timeline of Suessiales divergence from the rest of dinoflagellates. Evolutionary timescale along the different eras highlighting key geological events relevant to coral evolution, aligning with Symbiodiniaceae divergence, is shown at the bottom. Mass of preserved corals and sponges represented by black line (left y-axis), low latitude (30°S-30°N) ocean temperature in red (right y-axis), ocean chemistry of Mg/Ca ratio showing aragonite (dark blue) vs calcite sea (light blue); data were sourced from earlier studies (*1, 2, 13-15, 135, 136*).

*E. voratum* is estimated to have diverged ∼147 MYA in the late Jurassic (*1*). We did not find evidence for genome streamlining in *E. voratum*, but instead found genomic hallmarks associated with earlier-branching free-living lineages external to Family Symbiodiniaceae, e.g., more IE-containing genes and larger intron size than in S1 and S2. Given that *Effrenium* is also the sole genus within the Family that is exclusively free-living, we postulate that this lineage has not been impacted by symbiogenesis, in contrast to S1 or S2. Since the lineage diverged from S1, global events such as the breakup of Pangea (*11, 66*) and the Cretaceous-Paleogene mass extinction have occurred, along with changes in coral reef biomass (Fig. 6). Without additional evidence from the fossil record or ancient DNA analysis (oldest evidence of Suessiales is *P. glacialis* from just 9,000 years ago (*67*)), we cannot explain why *Effrenium* retained a free-living lifestyle. *Effrenium* and most other Symbiodiniaceae lineages (including free-living species from *Symbiodinium*) are capable of forming endolithic relationships with bacteria, e.g., as calcified biofilms (*68, 69*), but why *Effrenium* cannot form endosymbiosis with a host is unknown.

At the estimated time when S2 lineages diversified (∼109 MYA), shallow-water corals were described to be multiserial, flatter, and more efficient at harvesting light (*65*). This is coincident with a rise in ocean temperature (i.e., “greenhouse Earth”, when no continental glaciers exited) and the switch in ocean chemistry from an aragonite sea to a calcite sea, which would have made it difficult for corals to build their aragonite skeletons (*14*) (Fig. 6). In contrast, bivalve rudists that could build aragonite or calcite shells (*70*) radiated and flourished (*13, 15*). These taxa likely harboured photosymbionts (*12*), presumably ancestral Symbiodiniaceae, given that extant Symbiodiniaceae (e.g., *S. tridacnidorum*) can inhabit modern bivalves (*71*). Although genomes of S2 exhibit lower GC content than those of S1, our results clearly indicate that both S1 and S2 underwent genome streamlining, which led to convergent evolution of protein families associated with functions relevant to forming a symbiotic association, such as cell signalling, apoptosis, and photosynthesis (Fig. 3). We posit that symbiogenesis drove genome evolution in Symbiodiniaceae and elicited common features such as pseudogenisation, lowered mRNA editing and intron contraction, but some features (such as LINE retention and GC content) were affected differently in earlier vs. later-branching symbiotic lineages.

## Discussion

Our results provide strong evidence for a phase of genome reduction that occurred in the Suessiales ancestor. Therefore, the Symbiodiniaceae have smaller genome sizes than most free-living dinoflagellates, not because of the coral symbiosis, but due to more ancient selective constraints. These results are consistent with (but do not prove) the appealing idea that symbiosis offered an “escape” from reduced functional capacity due to genome reduction during the early stages of Symbiodiniaceae evolution. Regardless, the mixture of obligate free-living to facultative lifestyles among extant Symbiodiniaceae has resulted in divergent paths of genome evolution. Our results demonstrate the retention of ancestral Symbiodiniaceae genome features in *E. voratum* (in contrast to symbiotic lineages) despite multiple emergences of symbiogenesis over the past 200 million years. These observations support the notion that evolution favoured a free-living lifestyle for *E. voratum* (and by extension the genus *Effrenium*), likely due to local selective pressures. Therefore, *Effrenium* presents a useful free-living outgroup for studying the structural and functional genome features of symbiotic Symbiodiniaceae, and the implications of these features on ecology and evolution, including but not limited to host specificity and the “facultativeness” of symbiotic associations.

## Materials and Methods

### E. voratum cultures

Cell cultures of *E. voratum* RCC1521, rt-383, and CCMP421 were provided by the LaJeunesse Lab in Pennsylvania State University, U.S.A. They were incubated using Daigo’s IMK medium at 25°C under light-dark cycles of 14:10 hours. RCC1521 and rt-383 were maintained at the University of Queensland, CCMP421 at Florida International University.

### Microscopy

Cell cultures of *B. minutum* Mf1.05b.01, *D. trenchii* CCMP2556, *E. voratum* CCMP421, *S. microadriaticum* Cass KB8, and *Symbiodinium linucheae* SSA01 were grown in f/2 medium (*72, 73*) prepared using artificial seawater. The cells were fixed for 30 minutes in 4% v/v formaldehyde:f/2 medium (salinity 35 ppt). Cells were then pelleted by centrifugation (300 *g*, 5 min, room temperature [RT]) and washed twice using the f/2medium. All micrographs were captured at 100x (with a 0.63x adapter) on an Olympus BX 63 in brightfield mode.

### Extraction of genomic DNA

To extract genomic DNA (gDNA) for RCC1521 and rt-383, cells were pelleted by spinning the culture at 300 *g* (5 min, RT). Then the cell pellet was resuspended in 100-500 μL (higher volumes for more cells) prewarmed (at 60°C) lysis buffer (100mM Tris-HCl, 20mM EDTA, 4% CTAB (w/v), 1.4 NaCl, 1% 360,000 g·mol^-1^ PVP (w/v), 2% β-mercaptoethanol). This mixture was transferred to liquid nitrogen pre-chilled mortar pestle and ground to a fine powder and high molecular weight (HMW) gDNA was extracted as described in Rai (*74*). Briefly, Proteinase K (200 μL of stock at 20 mg/mL) was added, multiple rounds of chloroform:isoamyl alcohol (24:1 v/v) extractions were performed, the CTAB-DNA pellet was washed using ethanol, RNase A (4 μL of stock at 20 mg/mL) was added, and HMW gDNA was captured by incubating with 7.5 M ammonium acetate. The final HMW gDNA was resuspended in Tris-HCl (10 mM, pH 8) prewarmed at 50 °C, then was stored at -20°C until sequencing.

*E. voratum* CCMP421 cells were pelleted and snap frozen in liquid nitrogen and ground along with glass beads (diameter 425-600 µm), and the 2×CTAB method described in Stephens et al. (*19*) was followed. Briefly, the ground powder was transferred into a lysis buffer (100 mM Tris-HCl pH 8, 20 mM EDTA pH 8, 1.4 M NaCl, 2% w/v CTAB), mixed with RNAse A (final concentration 20 μg/mL) and incubated at 37°C (30 min), and then with Proteinase K (final concentration 120 μg/mL) at 65°C (2h). The standard extractions using phenol-chloroform-isoamyl alcohol (25:24:1 v/v; centrifuged at 14,000 *g*, 5 min, RT) and chloroform-isoamyl alcohol (24:1 v/v; centrifuged at 14,000 *g*, 5 min, RT) were then performed. The DNA was precipitated using chilled isopropanol (18,000 *g*, 5 min, 4°C), washed using chilled 70% ethanol, then stored in Tris-HCl (10 mM, pH 8) until sequencing.

### Extraction of total RNA

To extract total RNA from *E. voratum* RCC1521 for Iso-Seq sequencing, cell pellets were lysed via five freeze-thaw cycles with 425-600 µm diameter glass beads added to cell pellet, dipped in liquid nitrogen, vortexed, and thawed at 37°C), and then the QIAGEN RNeasy kit was used following the plant tissue protocol. To increase transcriptome diversity, we extracted more RNA using a different method. Total RNA was extracted for RNA-Seq (RCC1521 and rt-383) and Iso-Seq (from rt-383) following Acosta-Maspons et al. (*75*) with slight modifications. Cells were first pelleted by centrifugation (300 *g*, 5 min). The cell pellet was suspended in 100–500 μL lysis buffer (100mM Tris-HCl, 25 mM EDTA, 2% CTAB w/v, 2M NaCl, 0.75 g/L spermidine trihydrochloride, 4% β-mercaptoethanol). This mixture was transferred to a mortar and pestle that were prechilled in liquid nitrogen, and ground to a fine powder. The powder was swirled with liquid nitrogen and transferred to a Falcon tube (15 mL) standing in dry ice, then the liquid nitrogen was allowed to evaporate. Lysis buffer (5 mL; prewarmed at 65°C) was added, mixed by vortexing, and the tube was kept on ice for subsequent steps. An equal volume of chloroform:isoamyl alcohol (24:1 v/v) was mixed in; the mixture was divided into aliquots (2mL) in Eppendorf tubes and centrifuged (10,000 *g*, 10 min, 4°C). The clear supernatants were transferred to new tubes (2mL each) before the extraction with chloroform:isoamyl alcohol (24:1 v/v) was repeated. Nucleic acids were precipitated using LiCl (final concentration 2M) overnight at 4°C, and centrifuged (17,000 *g*, 30 min, 4°C). The resulting pellet was resuspended in the SSTE buffer (50 μL; 1 M NaCl, 0.5% SDS w/v, 10mM Tris-HCl pH 8, 1mM EDTA; prewarmed at 65°C); the samples were not set on ice to avoid undesirable precipitates. Further chloroform:isoamyl alcohol (24:1 v/v) extraction was performed (centrifuged 10,000 *g*, 10 min, 4°C), and the RNA pellet was precipitated using an equal volume of 2-propanol (incubated 10 min at RT, then centrifuged at 17,000 *g*, 10 min, 4°C). The pellet was washed with 80% ethanol (500 μL), dislodged by pulse vortexing (2 s), and the tube centrifuged (17,000 *g*, 5 min, 4°C); this step was repeated. The final pellet was air-dried (10 min) in a fume hood, resuspended nuclease-free water (25 μL), and stored at -80°C.

### Generation of genome data

For short-read sequencing, the libraries for RCC1521 and rt-383 were prepared using the Illumina TruSeq Nano kit with 350 bp targeted inserts following standard protocol, and then sequenced on the NovaSeq 6000 platform at Australian Genome Research Facility (Melbourne, Australia). For CCMP421, the genomic DNA library was prepared using the Chromium Genome Reagent Kit v2 Chemistry following the manufacturer’s protocol (Step 2 GEM generation and barcoding onwards) and then sequenced on the NovaSeq 6000 platform at Florida International University.

For Nanopore long-read sequencing of RCC1521 and rt-383, the ligation kit SQK LSK-109 was used following standard protocol. Each library was sequenced on a MinION flow cell and rt-383 gDNA was further sequenced using a PromethION flow cell at the Genome Innovation Hub (GIH), the University of Queensland (Brisbane, Australia). The sequence reads were base-called using guppy v4.0.11 as part of the MinKNOW software v20.06.18, with the minimum read quality filter of 7.

For PacBio long-read sequencing of RCC1521 and rt-383, the gDNA of RCC1521 (unsheared, for continuous long-read [CLR] library) and rt-383 (sheared in 15-20 Kb fragments with Pippin Prep (Sage Science), for HiFi library) were used for library preparation using the SMRTbell Express Template Prep Kit 2.0 following the manufacturer’s protocol.

The RCC1521 and the rt-383 libraries were sequenced on the PacBio Sequel II platform, respectively at the University of Washington PacBio Sequencing Services (Settle, WA, USA) and at the University of Queensland Sequencing Facility (Brisbane, Australia). The CLR reads for RCC1521 were acquired using the PacBio BAM2fastx toolkit, whereas the HiFi CCS reads for rt-383 were obtained using the CCS module of the SMRT Link pipeline v8.0.

### Generation of transcriptome data

Transcriptome data were generated for RCC1521 and rt-383. Illumina RNA-Seq libraries were generated using polyA-selection with the Dynabeads mRNA purification Kit and the Illumina Stranded mRNA Prep following standard protocols. Sequencing was performed on the Illumina NovaSeq 6000 platform at the Australian Genome Research Facility (Melbourne, Australia). PacBio Iso-Seq libraries were prepared using the NEBNext® Single Cell/Low Input cDNA Synthesis and Amplification Module (New England BioLabs) and the SMRTbell Express Template Prep Kit 2.0 following standard protocol and sequenced on the PacBio Sequel II at the University of Queensland Sequencing Facility.

### Transcriptome assembly and processing

For RNA-Seq data from RCC1521 and rt-383, adapters and unique molecular identifiers were removed using Illumina’s bcl2fastq v2.20.0.422, and the reads were trimmed (for polyG and polyA) and filtered using fastp v0.20.0 (-*A -L 35 -g -x --cut_front --cut_window_size 4 -- cut_mean_quality 15*). The processed reads were assembled using Trinity v2.9.1 (*76*) in both “de novo” (*--SS_lib_type RF --trimmomatic*) and “genome-guided” modes; for the latter, RNA-Seq reads were first mapped to the assembled genome using HISAT2 (*77*) before Trinity was run using *--SS_lib_type RF --genome_guided_bam --genome_guided_max_intron 70000*. We assessed the completeness of the each transcript set using BUSCO v5.1.2 (*78*) against the alveolata_odb10 database (Table S1).

Iso-Seq transcripts do not require assembly. The raw Iso-Seq sequences of RCC1521 and rt-383 underwent CCS generation and demultiplexing using the standalone modules CCS v4.2.0 and Lima v1.11.0. The rest of the IsoSeq processing steps (i.e., refining, clustering isoforms, and polishing) were conducted using the IsoSeq pipeline v3.3.0, resulting in a final set of high-quality transcripts.

### Estimation of genome size from sequencing data

Illumina short reads were used for estimating genome size based on *k*-mers. First, the reads from each genome dataset were processed to remove potential adapters (for 10X linked-reads of CCMP421, specifically the first 23 bases) and polyG tails using fastp v0.20.0 (*79*).

Jellyfish v2.3.0 (*80*) was used to obtain *k*-mers of sizes 17-31. The *k*-mer distribution was then plotted for each *k,* and the genome size was estimated as the sum of observed *k*-mers divided by the mean coverage (corresponding to the peak of the curve), averaged from the different *k*-mer sizes (Table S2). The ploidy of the genome datasets was assessed using GenomeScope2 (*81*) based on 21-mer distribution and a better fit was observed using a haploid (p=1) model in all three cases. Genome size estimations of other dinoflagellates (Table S12) were taken from Rizzo et al. (*82*), Hou and Lin (*28*), Sano and Kato (*83*), and Kohli et al. (*84*).

### *De novo* genome assembly

*De novo* genome assemblies were generated for RCC1521 and rt-383 adopting a hybrid-data approach, combining Illumina (short reads), PacBio (long reads), and Nanopore (long reads) data using MaSuRCA (*85, 86*) v4.0.1 for RCC1521, and v3.4.2 for rt-383, with the built-in CABOG as the final assembler; the key distinction between these two versions of MaSuRCA is the six-fold decrease in run-time in v4.0.1 relative to that for v3.4.2, with negligible impact on the yielded assemblies. For CCMP421, the *de novo* genome assembly was generated from 10X Genomics linked-read sequencing data using Supernova v2.1.1 (*87*).

RCC1521 and rt-383 assemblies were further scaffolded with L_RNA_scaffolder (*88*), using IsoSeq transcripts and *de novo* assembled transcripts from RNA-Seq (above). For CCMP421, linked-read distance information was first used to refine the assembly with ARBitR v0.2 (*-m 27k -s 10k*) (*89*) prior to scaffolding with L_RNA_scaffolder (*88*). Due to the low quality of the publicly available transcriptome data of CCMP421 (i.e., MMETSP1110 (*90*) with only 54% mapped to the corresponding assembled genome; Table S13), we used the *de novo* assembled transcripts from RCC1521 and rt-383 to scaffold the CCMP421 genome assembly via L_RNA_scaffolder.

To ensure high quality of each genome assembly, we identified and removed potential contaminant sequences of bacterial or archaeal sources following a decision tree based on analysis using BlobTools v1.1 (*91*) as described in earlier studies of algal genomes (*22, 92*); this step yielded the final assembly for each isolate. For each assembly, we assessed data completeness using BUSCO v5.1.2 (*78*) against the alveolata_odb10 database. Genome-sequence similarity among the three *E. voratum* isolates was assessed using nucmer implemented in the MUMmer package v4.0.0beta2 (*--mum*) (*93*).

### Identification of mitochondrial and plastid genome sequences

We identified putative mitochondrial and plastid genome sequences from the three isolates of *E. voratum*. To search for mitochondrial scaffolds, we followed Stephens et al. (*19*) by adopting BLASTn v2.10.0+ (*94*) search against the assembled genomes, using the protein-coding sequences of *B. minutum* mitochondrial genes as queries (GenBank accessions LC002801.1 and LC002802.1; *E* ≤ 10^-10^). We used BEDtools merge (*95*) to merge overlapping BLAST hits of the same gene (*-s -o collapse -c 1,2,3,4,5,6*), then used BEDtools intersect (*-wa -wb*) to check for overlaps with the predicted gene annotations. Genome scaffolds with BLAST hits that did not have any other predicted genes were considered putative mitochondrial genomes. We manually annotated the genes on the putative genome scaffolds using Artemis (*96*) with translation table 4.

To identify plastid genome fragments that are known to be shorter and may not be recovered in a hybrid genome assembly combing long- and short-read sequence data, we performed an independent short-read only genome assembly for each isolate using CLC Genomics Workbench v21.0.4 (Table S14). We used BLASTn search (*E* ≤ 10^-10^) using protein-coding sequences of plastid-encoded genes for *Cladocopium* sp. C3 (GenBank accessions HG515015.1-HG515028.1) as query, and annotated the putative genome scaffolds in Artemis (translation table 11). To look for empty minicircles, we first determined the core region of all plastid genome scaffolds. We masked coding regions of the scaffolds using BEDtools *maskfasta* and did pairwise BLASTn searches. The region that was common to most scaffolds was considered the core region. We then used this as query in BLASTn searches among unannotated *E. voratum* short-read genome scaffolds. A scaffold was considered empty if they did not have any hits in the NCBI nr database (November 2020).

Finally, we looked for evidence of circularisation in all of the recovered organellar genome scaffolds using nucmer (*--mum -l 0*) and mummerplot (*--layout -- png*) from MUMmer 4.0.0beta2 (*93*) to self-align the genome sequences and visualise the alignments.

### *Ab initio* prediction of protein-coding genes

To predict protein-coding genes, we used an integrated, multi-method workflow customised for dinoflagellates (*22*) incorporating transcriptome and protein evidence (pipeline available at https://github.com/TimothyStephens/Dinoflagellate_Annotation_Workflow). We first predicted repetitive elements from the genome assembly, followed by gene predictions based on a) the genome only, b) protein sequence similarity, and c) mRNA transcripts.

*De novo* repeat families were predicted from the genome assembly using RepeatModeler v2.0.1 (*97*), and these repeats were added to the Dfam database (dfam.org; downloaded June 2019) to guide RepeatMasker v4.1.0 (*98*) in repeat-masking the genome assembly. Next, GeneMark-ES v4.65 (*99*) was used for *ab initio* gene prediction on the masked genome assembly. Protein-based gene prediction was performed using MAKER v2.31.10 (*protein2genome*) (*100*) modified to recognise dinoflagellate alternative splice sites. It integrated the custom repeat library from the repeat analysis step, and BLASTn/x (*101*) searches were performed against the combined protein sequence database of SwissProt (released March 2020) and the Suessiales sequences, hereby “Suessiales_pep”, listed in Table S7 of Chen et al. (*22*).

For transcript-based gene prediction, Iso-Seq transcripts where available (i.e., for RCC1521 and rt-383) were mapped on the corresponding genome assembly using minimap2 v2.18 (*102*) for which the code was modified to recognise dinoflagellate alternative splice sites, using options *--secondary=no -ax splice:hq -uf --splice-flank=no.* The assembled transcripts from RNA-Seq (both *de novo* and genome-guided) for each isolate were mapped to the corresponding genome assembly using BLAT (*103*); for CCMP421, *de novo* assembled transcripts from RCC1521 and rt-383, plus the assembly of these reads guided by the CCMP421 genome, were used in this step (due to poor quality of the publicly available CCMP421 transcriptome data; see Tables S1 and S13). The resulting GFF3 files were input into PASA v2.4.1 (*--IMPORT_CUSTOM_ALIGNMENTS_GFF3 -- transcribed_is_aligned_orient -C -R --MAX_INTRON_LENGTH 70000*) (*104*) which was modified to recognise dinoflagellate alternative splice sites.

The PASA-predicted genes were filtered in the following steps: they were searched against the combined RefSeq (release 98) and the Suessiales_pep database using BLASTp v2.2.26 (e-value < 10^-20^, both query and subject coverage > 80%) (*101*), putative transposon sequences were removed via running HHBLITS v3.3.0 (*105*) and TransposonPSI v1.0.0 (*106*) against the UniRef30_3030_03 database from Uniclust (uniclust.mmseqs.com) (*107*), redundant sequences were removed using CD-HIT v4.8.1 (*-c 0.75 -n 5*) (*108*), and the script *Prepare_golden_genes_for_predictors.pl* from the JAMg pipeline (https://github.com/genomecuration/JAMg) was used to produce a highly curated set of “golden genes”. These golden genes were used to guide the *ab initio* gene prediction tools SNAP (*109*) and AUGUSTUS v3.4.0 (*110*) on the repeat-masked genome assembly. Gene models from the five tools (MAKER, GeneMark-ES, PASA, SNAP, AUGUSTUS) were integrated using EVidenceModeler v1.1.1 (*111*). Finally, the resulting gene models were refined to correct exon boundaries, add UTRs, and incorporate gene models from alternative splicing using the *Load_Current_Gene_Annotations.dbi* and *Launch_PASA_pipeline.pl* from the PASA pipeline (*112, 113*) iteratively for three rounds to yield the final gene models.

We assessed the completeness of the predicted protein sequences using BUSCO v5.1.2 (*78*) against the alveolata_odb10 database in “protein” mode. Genes with transcript support were identified by aligning transcripts for each isolate to their predicted coding sequences using BLASTn (e-value < 10^-5^, percent identity ≥ 90 %, subject cover ≥ 50 %).

### Functional annotation of predicted genes

Functions of the predicted protein sequences were annotated based on BLASTp searches (e-value < 10^-5^ and query/subject cover ≥ 50 %) against the SwissProt (2022_01) database.

Then, those that had no hits were searched against TrEMBL (2022_01) database; the function of the top protein hit was assumed to the putative function of the query protein. The UniProt IDs were converted to Gene Ontology (GO) terms via the UniProtKB ID mapping tool (https://www.uniprot.org/id-mapping) in December 2022.

### Inferring phylogenetic tree based on multiple sequence alignment

To infer phylogenies for the 18S rDNA and ITS2 marker sequences, we first recovered 18S and ITS2 sequences from the three genome assemblies of *E. voratum* using BLASTn. Reference sequences for the 18S rDNA were downloaded from https://doi.org/10.5061/dryad.1717129 (79 sequences) (*1*) and ITS2 from SymPortal (https://symportal.org; “published post-MED sequences” of 8,409 sequences downloaded 10 September 2021) (*114*), respectively. For each marker sequence set, multiple sequence alignment was generated using MAFFT v7.471 in *mafft-linsi* mode (*115*), trimmed using trimAl v1.4.rev15 *(-automated1*) (*116*), from which a maximum-likelihood phylogenetic tree was inferred using IQ-TREE v2.1.3 *(-nm 2000 -bb 2000 -m MFP*) (*117*). The 18S rDNA tree based on LaJeunesse et al. (*1*) and the associated divergence times was overlayed with major geological events and visualised using tvBOT (*118*).

To reconstruct a reference species tree of dinoflagellates based on strictly orthologous protein sequences, we incorporated 1,603,073 predicted protein sequences from 33 dinoflagellate taxa, comprising 21 Suessiales taxa (including the three *E. voratum* isolates in this study) and 12 other taxa external to Suessiales (Table S12) (*90, 119, 120*). These sequences were clustered into homologous sets using OrthoFinder v2.5.4 (*121*), from which a species tree was estimated from strictly orthologous sets.

### Alignment-free phylogenetic inference and core *k*-mers

We used an alignment-free approach to infer phylogenetic relationships from (a) whole-genome sequences (WGS) and from distinct genomic regions of (b) repeat-masked WGS, (c) coding sequences (CDS), (d) introns, (e) annotated repeats, and (f) predicted protein sequences. Each of these distinct regions were extracted from assembled genome sequences using *gff3_file_to_feature_files.pl* implemented in PASA (*104*). We followed Lo et al. (*39*) to identify optimal *k*-mer length (*k*) for each of these datasets. Briefly, for each dataset, *k*-mers at varied length *k* were enumerated using Jellyfish v2.3.0 (*80*); for all datasets, odd-numbered *k* between 13 and 27 were used, except for the repeats dataset (odd-numbered *k* values between 13 and 51 were used) and protein sequences (odd-numbered *k* values between 3 and 9 were used). For each dataset except the protein sequences, the optimal *k* was determined based on the cumulative proportion of unique *k*-mers and the cumulative proportion of distinct *k-*mers, at the point when distributions of both proportions reached a plateau (Fig. S7); *k* value determined this way was found to yield the greatest distinguishing power for phylogenetic analysis (*122*). For protein sequences, we inferred alignment-free phylogenies (see below) from each *k* and chose the *k* with a topology that best matched the 18S rDNA alignment-based tree as implemented in Lo et al. (*39*). The optimal *k* was identified as 23 for WGS and repeat-masked WGS, 19 for CDS, 21 for introns, 51 for annotated repeats, and 9 for protein sequences. Jellyfish v2.3.0 was used to extract *k-*mers at the corresponding optimal *k* length for each dataset, with the option *-C* used to enforce strand-specific directionality for the intron and CDS datasets.

To infer alignment-free (AF) phylogenies based on *k*-mers (i.e. using the optimal *k* for each corresponding dataset identified above), we derived pairwise distance based on 𝐷^*S*^_2_ statistic (*123*) following Chan et al. (*124*), using *d2ssect* (https://github.com/bakeronit/d2ssect). These pairwise distances were then used to infer a phylogenetic tree using *neighbor* implemented in PHYLIP v3.698 (*125*). For each tree, we assessed node support based on jackknife analysis of 100 “pseudo-replicates” following Bernard et al. (*126*). In each pseudo-replicate, 40% of the data, in iteratively subsampled 100-bp regions at random, were deleted using the Python script *jackknife.py* from which an AF tree was inferred; the R script *Jackknife.r* was then used to calculate jackknife support value in percentage, among the pseudo-replicate trees, for each node in the original AF tree. These scripts are available at https://github.com/chanlab-genomics/alignment-free-tools.

To identify core *k-*mers (*127*) that are shared by all 21 Suessiales genomes used in this study, we used the optimal *k* = 23 for the WGS dataset. Using the extracted 23-mers from the entire WGS dataset as input, core 23-mers were identified using the bash command *comm* (−12). BEDtools (*95*) *intersect* was used to find regions of overlap between the core *k-*mers and different genomic features.

### Analysis of gene family evolution

To examine the gene family evolution between *E. voratum* and the earlier-/later-branching symbiotic lineages of Symbiodiniaceae, we grouped 21 Suessiales protein sequence datasets into three groups: Ev, S1, and S2 (Table S3), with Po (*P. glacialis* CCMP1383 and CCMP2088) as the outgroup. The 811,661 protein sequences from the 21 Suessiales taxa were clustered into homologous families using OrthoFinder v2.5.4 (*121*). We then subset the shared/exclusive protein families among the different groups (Ev, S1, S2, and Po). GO enrichment was performed using the topGO package in R (algorithm = “elim”, statistic = “fisher”) for six comparisons: (a) shared genes in Ev+Po (test set) versus all genes in Ev+Po (background), (b) shared genes in S1+S2 versus all genes in S1 and S2, (c) shared genes in S1+S2+Po versus all genes in S1, S2, and Po, (d) shared genes in S1+S2+Ev+Po versus all genes in the 21 taxa, (e) exclusive genes to S1 versus all S1 genes, and (f) exclusive genes to Ev versus all Ev genes.

### Identification of pseudogenes

Pseudogenes were identified following the method described in González-Pech et al. (*21*) based on tBLASTn search using the predicted protein sequences as query against the corresponding genome sequences for which the predicted gene model sequences were masked. Matched regions (≥ 75% identity) were considered fragments of pseudogenes, and fragments at no more than 1 Kb apart and in the same orientation were considered collectively as a pseudogene.

In this analysis, we focused on 752,954 protein sequences from the 19 Suessiales taxa, specifically excluding *S. natans* and *S. pilosum* from S1 to avoid signatures of free-living lifestyle in these taxa interfering with potential signatures of symbiogenesis. The protein sequences were first clustered into homologous families using OrthoFinder v2.5.4 (*121*). We define the extent of pseudogenisation, \J¢, as the ratio of the number of putative pseudogenes to the number of putative functional genes in a homologous family. We determined this value independently for Ev (ψ_Ev_) against that for S1 (ψ_S1_), S2 (ψ_S2_), and the combined S1 and S2 (ψ_S1+S2_); a protein family with ψ_S1_ > ψ_Ev_ indicates a greater extent of pseudogenisation in S1 than in Ev. We then ran a one-way ANOVA test and Tukey’s test using the R package *rstatix* (https://cran.r-project.org/package=rstatix) to assess correlation among the Ψ values of Ev, S1, S2, and the combined S1+S2 groups.

To assess the robustness of our approach to the inflation parameter (*I*) in single-linkage clustering within OrthoFinder that modulates granularity (i.e., sizes and numbers) of the resulting protein families, we ran OrthoFinder independently using *I* = 1.1, 1.3, 1.7 and 2.0 in addition to the default value of 1.5, and performed the analysis of pseudogenisation per above.

### Analysis of mRNA editing

For the analysis of mRNA editing, we focused on *E. voratum* RCC1521, and the representative genomes for S1 (*S. microadriaticum* CCMP2467) and S2 (*D. trenchii* CCMP2556), for which high-quality genome and transcriptome data are available. Editing of mRNAs was identified using JACUSA2 (*128*), based on observed nucleotide variants in the mapping of transcripts onto the genome, relative to the mapping of genome sequence reads onto the genome. First, gDNA reads were mapped on each genome using BWA-mem v0.7.17-r1198 (*129*) using default settings. Then, for each genome, RNA-Seq reads were mapped using HISAT2 v2.2.0 *(--rna-strandness RF*) (*77*) to the assembled genome sequences with a HGFM HISAT2 index (*hisat2-build --exon --ss*) informed by the annotated splice sites. To generate the index, the gene annotation file in GFF3 was converted to the GTF format using Gffread (*130*), and splice site and exon locations extracted with the *hisat2_extract_splice_sites.py* and *hisat2_extract_exons.py* scripts. Iso-Seq reads, where available, were mapped using minimap2 v2.18 (*102*) for which the code was modified to recognise alternative splice sites of dinoflagellates, with options *--splice-flank=no – secondary=no -ax splice:hq -uf –junc-bed*.

Duplicate mappings were removed from the gDNA BAM files using Picard MarkDuplicates (*ASSUME_SORTED=true REMOVE_DUPLICATES=true CREATE_INDEX=TRUE VALIDATION_STRINGENCY=LENIENT*). The MD field documenting mismatched and deleted bases was added to the gDNA BAM files with samtools calmd (*-b*), as required as input for JACUSA2. JACUSA2 analysis was performed on the gDNA, RNA-Seq, and Iso-Seq BAM files using option *-a D,Y,H* to remove false positives caused by read starts/ends, indels, splice sites, and homopolymers. Due to the different strand directionality, *-P2 RF-FIRSTSTRAND* was specified for the runs incorporating RNA-Seq data, and *-P2 FR-SECONDSTRAND* for the runs using Iso-Seq data. The results from JACUSA2 were overlayed using BEDtools intersect (*95*) at *-s -wo* with predicted genes (including isoforms), and the edited sites were visualised using JACUSA2helper (https://github.com/dieterich-lab/JACUSA2helper).

We followed Liew et al. (*56*) to assess 5′ bias in the location of RNA editing and the propensity for edits to occur together. We calculated the frequency of the locations of each edit with respect to the features they were in (exon, intron, gene), normalised by the length of each feature. For each edit, its distance (in bp) to the closest upstream edit, and that to the closest downstream edit where all within the same gene, were determined. The average of these two values was used as the per-edit observed distance to neighbouring edits.

### Analysis of introner elements

To identify introner elements (IEs), we used Pattern Locator (*131*) to search for the patterns described in Farhat et al. (*53*): inverted repeats of 8-20 nucleotides within 30 bases of the 5′ and 3′ ends of each intron, flanked by direct repeats of 3-5 nucleotides. We first used Seqkit (*132*) to obtain the first and last 30 bases at each end of introns, then used Pattern Locator to identify the IEs.

## Acknowledgements

We are grateful to Todd LaJeunesse and Hannah Reich who generously supplied the cell cultures of two *Effrenium voratum* strains (RCC1521 and rt-383) used in this study. This project is supported by high-performance computing facilities at the National Computational Infrastructure (NCI) National Facility systems through the NCI Merit Allocation Scheme (Project d85) awarded to CXC, the University of Queensland Research Computing Centre, and computing facility at the Australian Centre for Ecogenomics, School of Chemistry and Molecular Biosciences at the University of Queensland.

## Author contributions

Conceptualization, SS, KED, DB and CXC; methodology, SS, KED, YC, SKR, AJB, VM and CXC; formal analysis, SS, KED, YC, RL, GL, MDAF, and VM; investigation, SS, KED; resources, SS, SKR, AJB, MRL, CXC, writing—original draft preparation, SS; writing— review and editing, SS, KED, DB, and CXC; visualisation, SS; supervision, KED, DB, CXC; funding acquisition, MRL, DB and CXC. All authors have read and agreed to the published version of the manuscript.

## Data and materials availability

All sequencing data generated from this study are available on NCBI GenBank via BioProject accession PRJEB61191. The assembled genome, predicted gene models and proteins, and the identified organellar genome sequences are available at https://doi.org/10.48610/1f0377a.

