## Supplementary Figures S1-S7 for "Massive genome reduction occurred prior to the origin of coral algal symbionts"

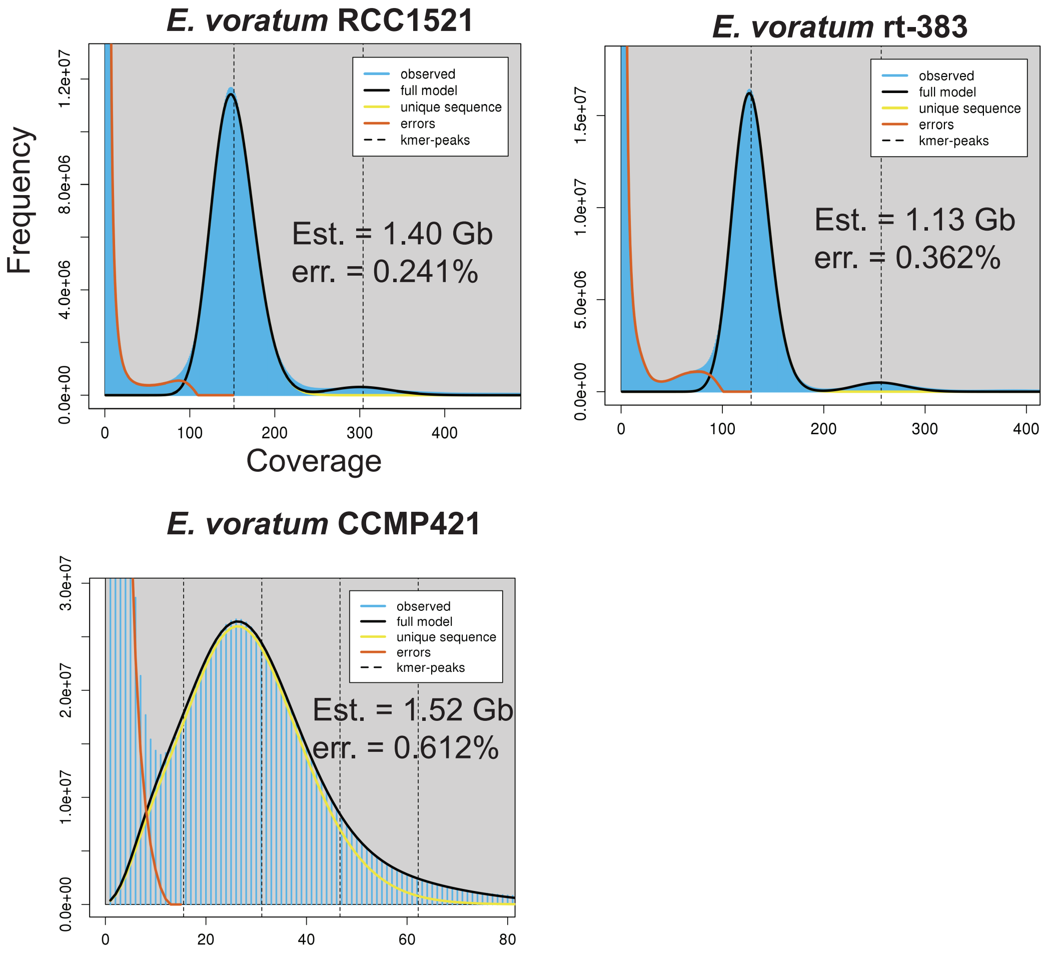


**Fig. S1.** GenomeScope profiles of 21-mers from gDNA reads of *E. voratum* showing the fit to a haploid model and its error rate, and the estimated haploid genome size.

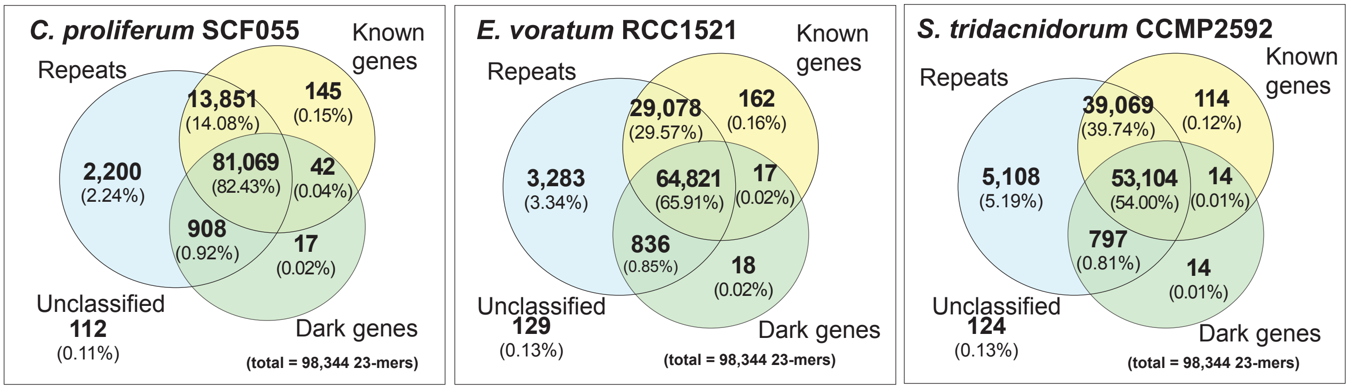


**Fig. S2.** The proportion of core *k*-mers among the 21 Suessiales taxa that correspond to different genomic regions of known genes (yellow), genes of unannotated functions (green), and repeats (blue) in *C. proliferum*, *E. voratum*, and *S. tridacnidorum*.

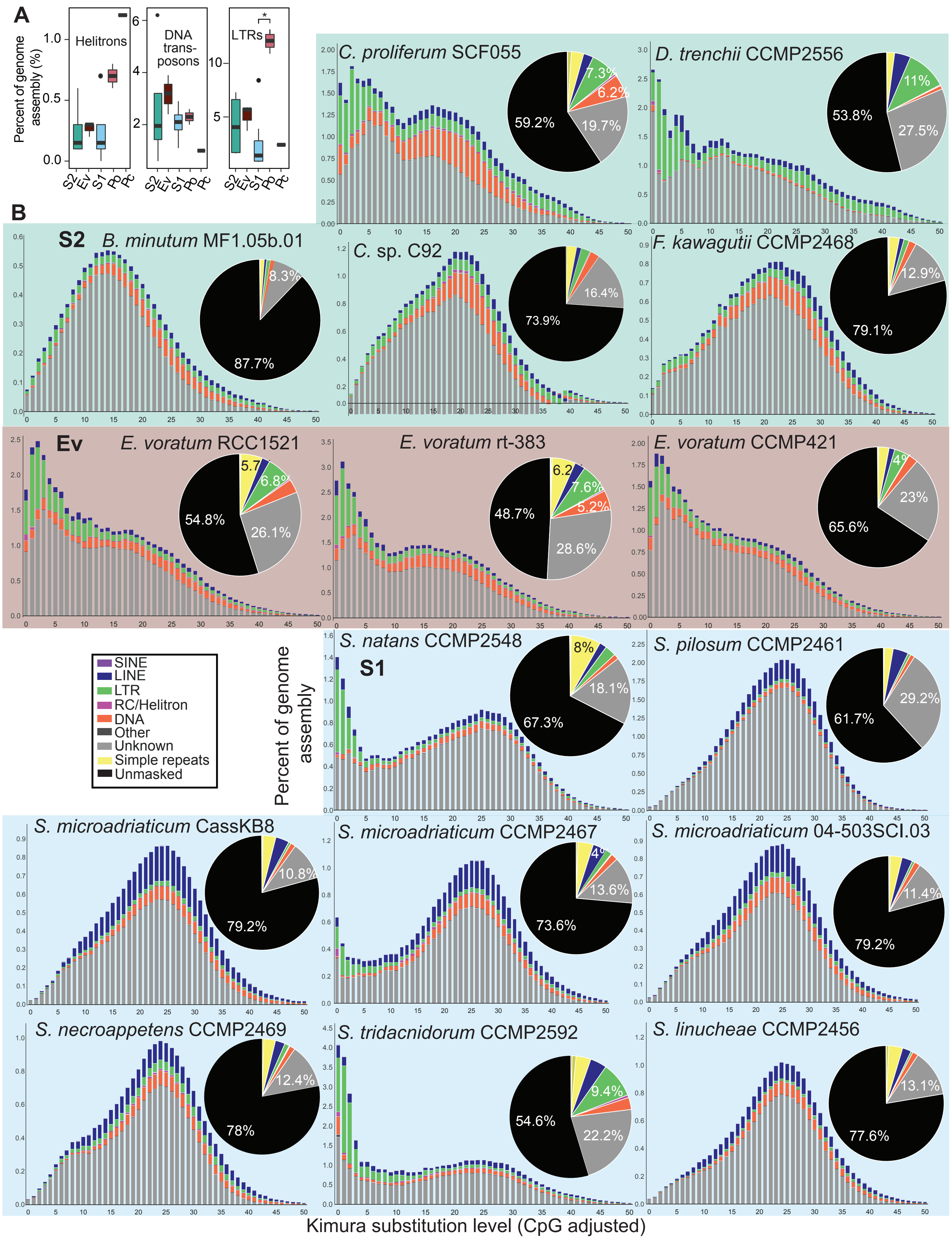


**Fig. S3.** Mobile elements in dinoflagellates, showing (A) proportions of helitrons, DNA transposons, and LTRs in S2, Ev, S1, Po, and *Prorocentrum cordatum* (Pc), and (B) repeat landscapes in genomes of Symbiodiniaceae, for S1, Ev and S2, showing the percentage of repeat types in each genome assembly and their level of conservation (Kimura substitution level).

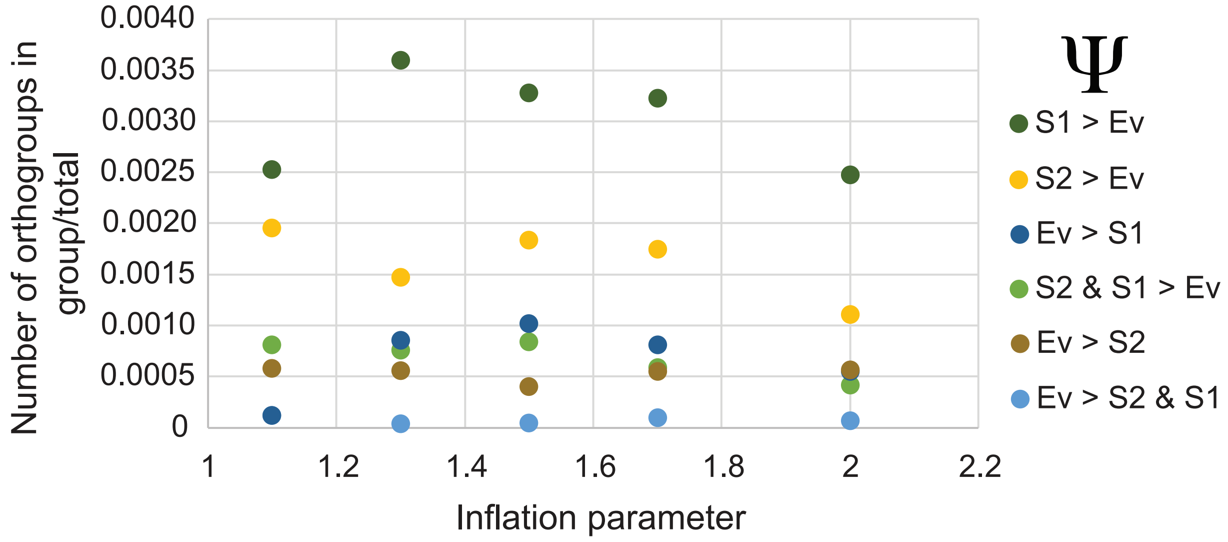


**Fig. S4.** The proportion of homologous groups in the distinct groupings of pseudogenisation analysis identified at varied inflation parameters implemented in OrthoFinder.

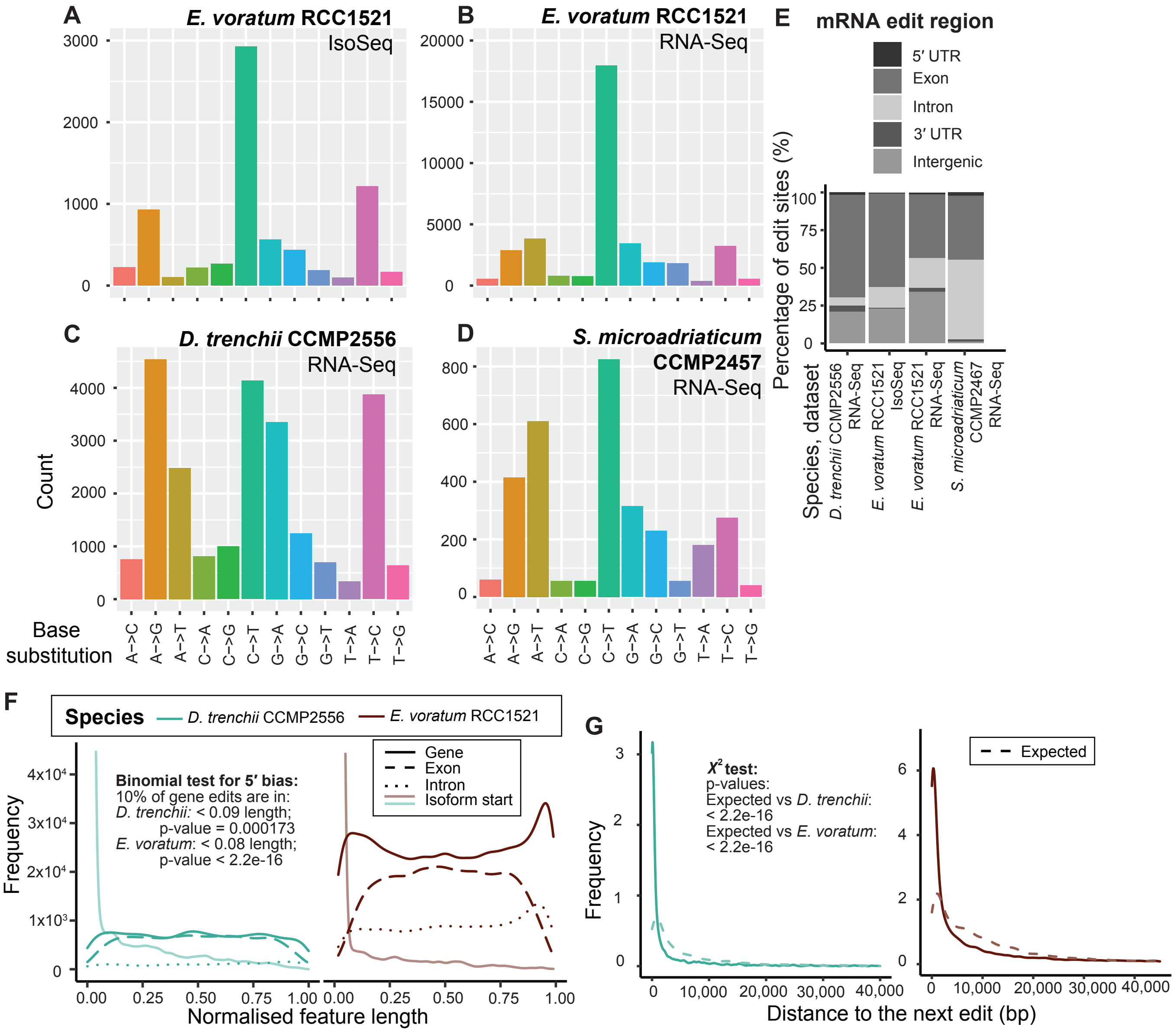


**Fig. S5.** Editing of RNAs in dinoflagellates showing different edit types identified using (A) *E. voratum* RCC1521 Iso-Seq, (B) *E. voratum* RNA-Seq, (C) *D. trenchii* CCMP2556 RNA-Seq, and (D) *S. microadriaticum* CCMP2457 RNA-Seq datasets, (E) locations of RNA edits, (F) frequency of RNA edits in *D. trenchii* CCMP2556 and in *E. voratum* RCC1521 along relative positions of genes, and (G) distances relative to the next edit within a gene.

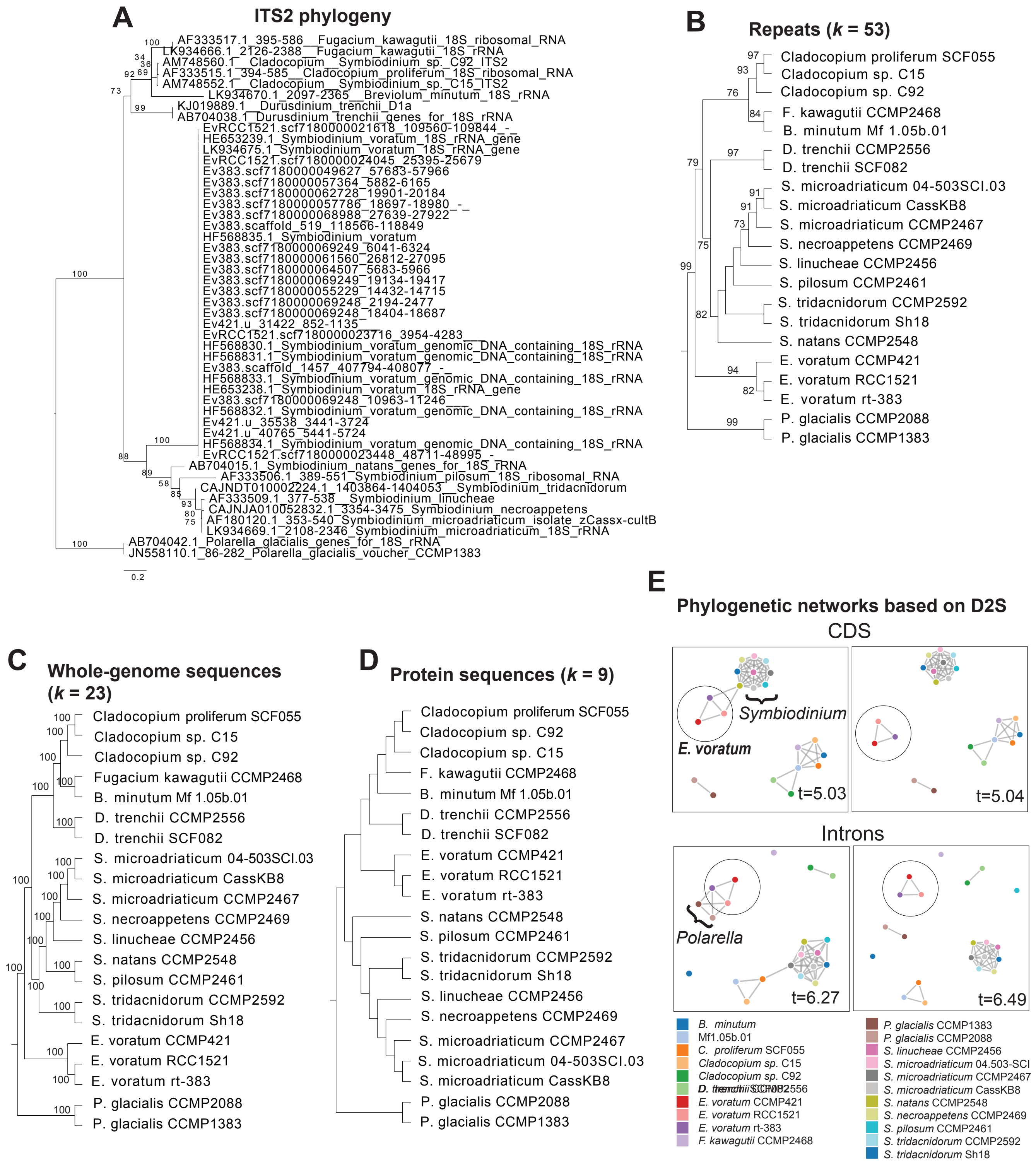


**Fig. S6.** Phylogenetic relationship of Suessiales inferred as trees based on (A) alignment of ITS2 sequences showing ultrafast bootstrap support value from 2,000 samples on each branch (unit in number of substitution per site); as tree topologies based on *k*-mers derived from (B) annotated repeat regions, (C), whole-genome sequences, and (D) protein sequences, showing jackknife support ≥ 70% from 100 pseudo-replicates where available; and as networks using (E) *k*-mer-derived distances from protein-coding and intron sequences, shown at distinct similarity thresholds t relative to split of *E. voratum* from other taxa.

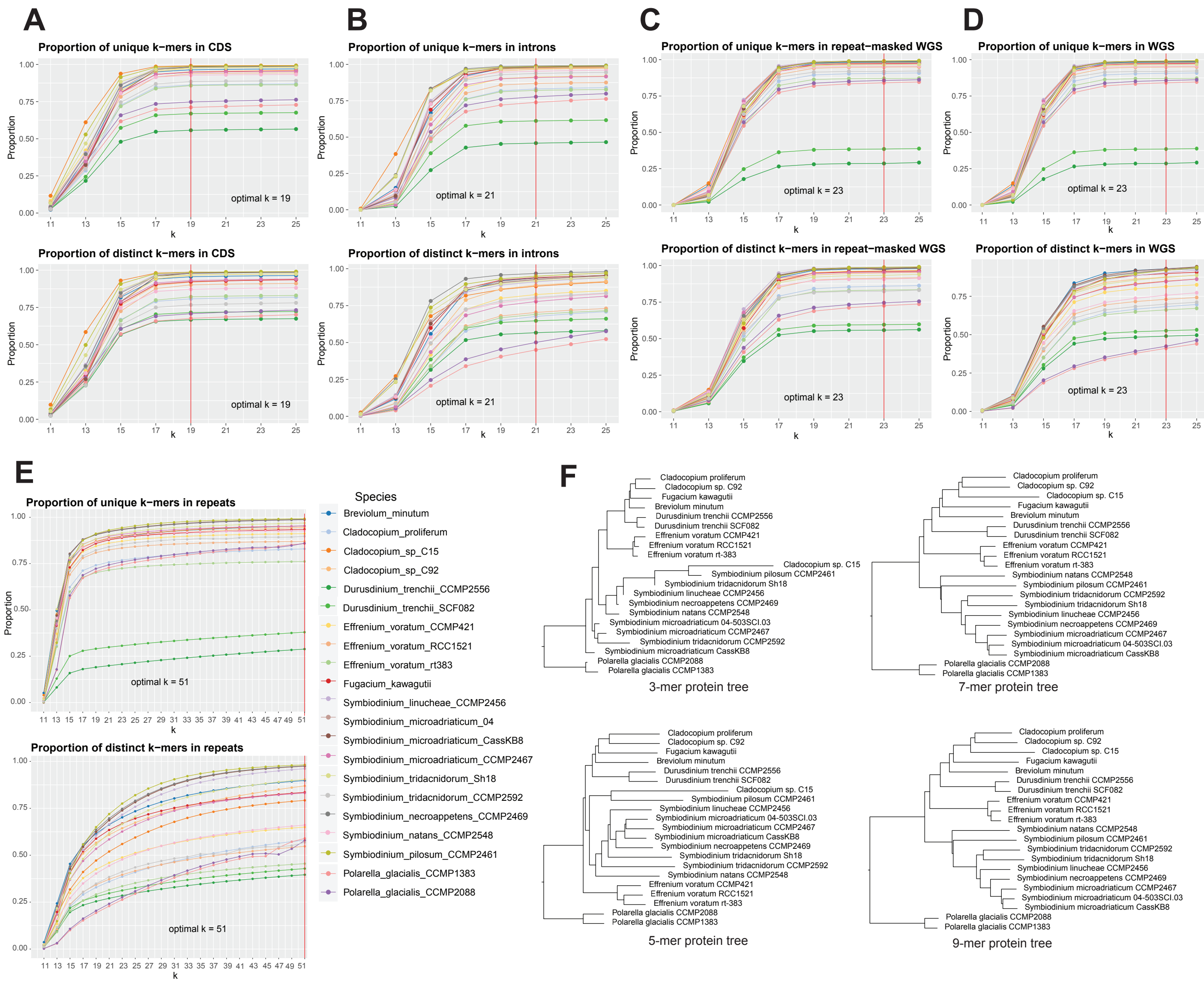


**Fig. S7.** Proportions of unique and distinct *k*-mers extracted from Suessiales shown for (A) CDS, (B) introns, (C) repeat-masked whole genome sequences, (D) whole genome sequences, and (E) repetitive regions, with the corresponding optimal *k* marked with a red line, and (F) the four protein trees inferred based on distinct values of *k*.
